# SQANTI-evidence: isoform-resolved, evidence-tiered structural annotation of unannotated genomes from long-read RNA-seq

**DOI:** 10.64898/2026.11.01.763455

**Authors:** Pablo Atienza López, Ana Conesa

## Abstract

Long-read RNA sequencing captures full-length transcripts, yet current annotation pipelines fail to properly curate the data for gene prediction. We present SQANTI-evidence, a novel software that annotates a genome from long reads alone, without the need for a reference annotation or protein database. SQANTI3-curated transcripts enter the annotation as observed, with isoforms and untranslated regions; AUGUSTUS predicts genes at loci without read support. In benchmarks, the curated layer was the most precise gene set in five of seven species, recovered twice as many exact isoforms as BRAKER in mammals and placed 3′ ends nearest the poly(A) site. We annotated 13 ERGA datasets.

## Background

Long-read sequencing technologies have rapidly expanded in recent years and have transformed the field of genomics [1,2]. Their main advantage over short-read technologies is their ability to resolve complex repetitive regions and long-range genomic structures. Long reads have greatly improved our capacity to assemble genomes, to the point that telomere-to-telomere assemblies are now within reach for many species [3]. As a consequence, the number of available reference genomes has grown exponentially, and initiatives such as the Earth BioGenome Project [4] and its European node, ERGA [5], are producing assemblies for species that have never had a high-quality reference genome.

This explosion of new genome assemblies creates a pressing need for genome annotation, the process by which genes and transcripts are assigned to defined genomic coordinates. Traditionally, genome annotation has used one of three approaches: *de novo*, *evidence-based* and *evidence-driven*. *De novo* approaches rely only on the genomic sequence to predict gene structure. They use previously known genes to train statistical models, such as hidden Markov models (HMMs) [6] or deep-learning architectures [7,8], that learn where gene boundaries lie and how gene structures are organized. These methods are effective and widely used, but have limited ability to identify genes whose structures differ from those in the training set. *Evidence-based* approaches rely solely on empirical evidence from multiple sources, such as RNA-seq or proteomics data [9,10]. They capture gene structures precisely, but are limited by the quality and depth of the available data. For example, a tissue- or condition-specific gene that is not expressed in the sequenced sample cannot be annotated. Finally, *evidence-driven* methodologies combine the strengths of both strategies. They predict gene structures from the genome sequence and use extrinsic evidence as hints to guide this prediction [11]. Evidence-driven methods can integrate different types of evidence to increase the scores of gene models that agree with the hints, or to penalize predictions not supported by them, without restricting the prediction to regions where evidence is present.

Transcriptomics data are among the most frequently used sources of evidence in annotation efforts. As with genome assembly, long-read sequencing has opened new possibilities for transcriptome analysis. Long-read RNA sequencing (lrRNA-seq) captures full-length transcripts and has expanded the reference transcriptomes of even well-annotated organisms [12]. However, lrRNA-seq data are not free of errors and artifacts, and benchmarking studies such as LRGASP have shown that the choice of analysis method strongly affects the transcript models inferred from long reads [13]. SQANTI3 is a widely used tool for the quality control and curation of long-read transcriptomes, which combines sequence features and orthogonal data to identify and remove artifactual transcript models [14,15].

Long-read transcripts can be used in *de novo* genome annotation, either as the sole source of evidence or as hints for evidence-driven methods. Several annotation tools accept long reads as input. EviAnn [16], LoReAn [17] and EGAPx [18] align them to the genome and build or select gene models from the alignments. BRAKER3 [19] in its long-read mode and its latest version, BRAKER4 [20], assemble them with StringTie2 [9], use the assembled transcripts as hints and apply GeneMark-ETP to infer protein-coding gene models. However, none of these tools curate the long-read data for the artifacts typical of lrRNA-seq libraries, none incorporates the observed isoforms into the annotation as such, and none records which models were observed in the sample rather than assembled or predicted. We have previously shown that curating long reads with SQANTI3 before prediction improves AUGUSTUS gene models [21]. However, this approach had several limitations. First, SQANTI3, designed primarily for annotated genomes, required a reference annotation for the curation step, which limited its applicability to new genomes. Second, the approach returned a single transcript per gene, thus failing to exploit the capacity of long reads to resolve alternative isoforms. Third, the method was tested only in mammalian species, and its performance in other clades remained unknown. Finally, the initial work delivered only a set of custom scripts, lacking the modularity, portability and reproducibility needed to support large-scale annotation efforts.

To address these limitations, we present SQANTI-evidence, a modular and highly customizable genome annotation workflow built on *ab initio* models combined with SQANTI3. SQANTI-evidence uses SQANTI3 and a set of filtering rules to curate the long-read transcriptome without a reference annotation, incorporates the curated transcripts into the annotation with their isoforms and untranslated regions (UTRs), and uses *de novo* gene prediction to cover the gene space missed by the transcriptomic data. We extensively benchmarked the workflow to understand the value of curating lrRNA-seq data for genome annotation. Each design decision was evaluated on eight model organisms, and the final annotations were compared with those of BRAKER3 and BRAKER4. Finally, we applied SQANTI-evidence to annotate newly assembled genomes from the ERGA pilot project for which lrRNA-seq data were available, and made these annotations publicly available through Zenodo. Our results show that long-read transcriptomes can be curated without a reference annotation and incorporated directly into genome annotations, providing gene models, isoforms and transcript ends that are observed rather than predicted.

## Results

### Overview of SQANTI-evidence and of the benchmark

SQANTI-evidence annotates a genome assembly from lrRNA-seq data alone, without requiring a reference annotation or an external protein database (Fig. 1). The workflow reconstructs transcript models from the reads with IsoQuant and curates them with SQANTI3 using only features intrinsic to each transcript. The curated transcripts, including their isoforms and untranslated regions (UTRs), are incorporated directly into the genome annotation to form the curated layer (Tier 1). To annotate genomic regions not covered by these transcripts, AUGUSTUS is trained on conserved BUSCO orthologs, supplemented with curated transcripts, and guided by hints derived from the curated transcripts and from their predicted proteins aligned back to the genome. Gene models predicted by AUGUSTUS at loci without curated transcripts form the predicted layer (Tier 2). Each model in the final annotation is labeled with its layer in a tier attribute, allowing users to distinguish transcript-supported models from computational predictions and to filter the annotation according to their needs.

**Figure 1.**
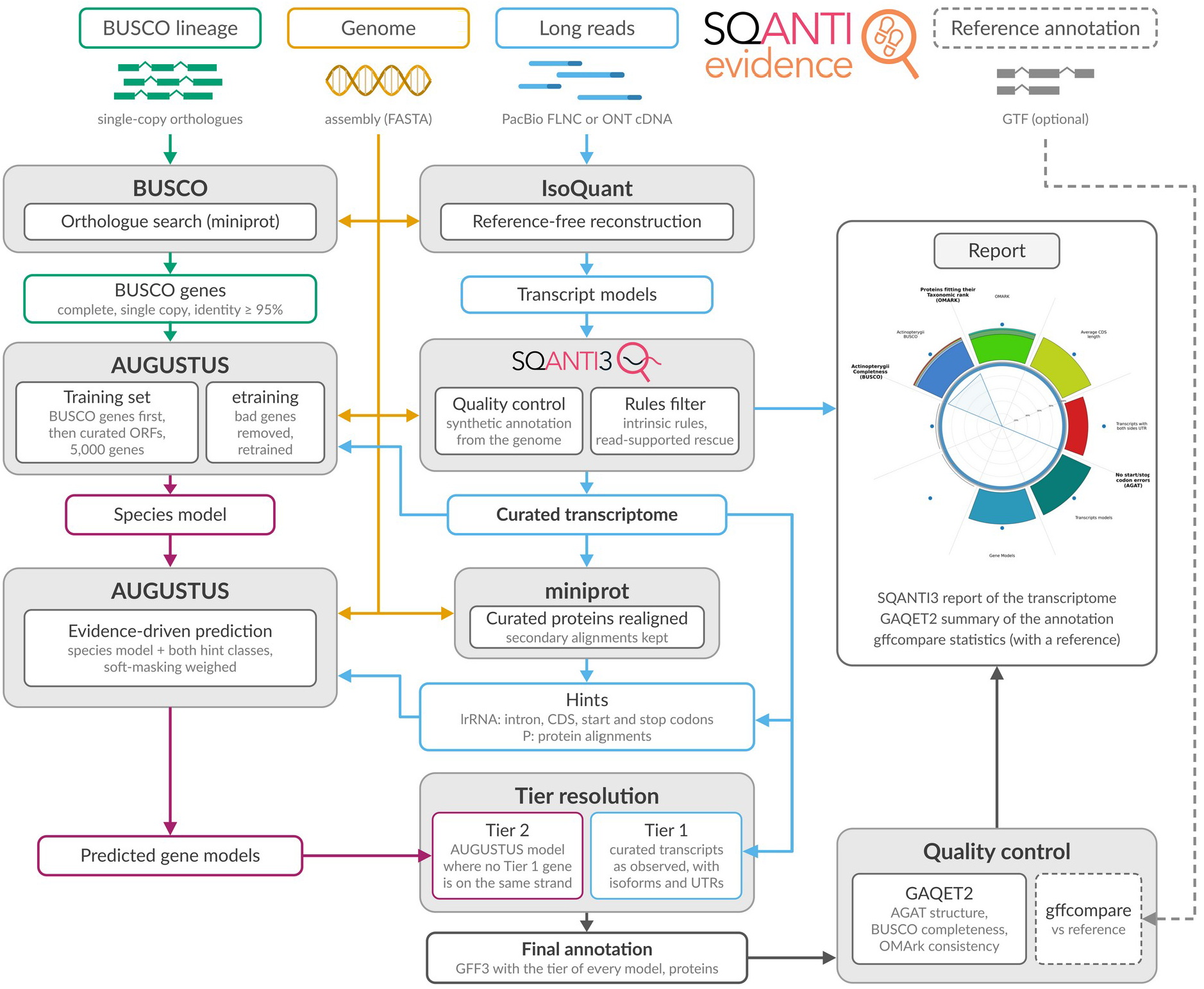
| The SQANTI-evidence workflow. Long reads (PacBio FLNC or ONT) and a genome assembly enter the workflow. IsoQuant reconstructs transcript models without a reference; SQANTI3 classifies them against a synthetic annotation, so every quality descriptor derives from the genome sequence, and the flexible filter keeps a model that passes all intrinsic tests or fails exactly one of the intra-priming, PSAURON or canonical-junction tests with ten or more full-length reads. The AUGUSTUS species model is trained on BUSCO orthologs placed by miniprot, completed with the dominant ORFs of the curated transcripts up to 5,000 genes, and retrained after removing the genes that etraining rejects. The curated proteins are realigned to the genome with miniprot, keeping secondary alignments as evidence for unexpressed paralogs. Intron, CDS and CDS start and stop hints are derived from the curated transcripts (source lrRNA); the protein alignments form a second hint class (source P). AUGUSTUS predicts in evidence-driven mode with both hint classes. Curated transcripts are written to the annotation exactly as observed (Tier 1, blue); an AUGUSTUS model is added only where no Tier 1 gene lies on the same strand (Tier 2, magenta), without an evidence test. GAQET2 and, when a reference is supplied, gffcompare score the result. Dashed lines, optional reference annotation, used only by gffcompare. Every feature in the output GFF3 carries a tier attribute.

We evaluated SQANTI-evidence in three stages. First, we assessed the main design choices of the pipeline using eight model organisms with well-curated reference annotations and publicly available long-read data, spanning genome sizes from 12 Mb to 3.3 Gb (Table 1). Second, we benchmarked the resulting annotations against those generated by BRAKER3 and BRAKER4 on the same species. Finally, we applied SQANTI-evidence to 13 datasets from the ERGA initiative, for which genome assemblies and long reads were available but reference annotations were generally lacking, with the exception of a *Caenorhabditis elegans* dataset used as a sequencing-depth control.

**Table 1.** | Species and datasets. Read counts: primary alignments of the FLNC reads of the final run (*Arabidopsis*: all FLNC reads in the input BAM). Error rate: mismatches plus indels per aligned base (samtools stats [49]), measured on about the first 400,000 primary alignments of each coordinate-sorted alignment file (*Arabidopsis*: a 200,000-read minimap2 spot check). IsoQuant models: transcript models reconstructed by IsoQuant before filtering including coding and non-coding. Table S1 and Results count the coding models only (e.g. 37,286 of 44,765 in human).

| **Species** | **Assembly** | **Size (Mb)** | **Reference** | **Long-read data** | **Platform** | **Reads** | **Error rate** | **IsoQuant models** | **BUSCO lineage (training)** |
|---|---|---|---|---|---|---|---|---|---|
| *S. cerevisiae* S288C | R64-1-1 | 12.2 | Ensembl 115 | PRJNA883530 (SRR21679915, SRR23265078, SRR21679893, SRR23265077), 4 runs pooled | Sequel II Iso-Seq | 991,895 | 1.03% | 5,008 | saccharomycetaceae_odb12 |
| *C. elegans* N2 | WBcel235 | 100.3 | Ensembl 115 | PRJEB121150 (ERR17585088–90), 3 replicates | Revio Kinnex FLNC | 18,125,019 | 0.12% | 20,417 | nematoda_odb12 |
| *A. thaliana* (“Bön 1”) | TAIR10 | 119.7 | Ensembl Plants 62 | ERR15191386 | Sequel II Iso-Seq | 722,673 | 0.65% | 17,241 | brassicales_odb12 |
| *D. melanogaster* | BDGP6.54 | 143.7 | Ensembl 115 | SRR29784392, SRR29784417, SRR29784414, merged | PromethION cDNA | 6,225,915 | 6.19% | 10,108 | drosophila_odb12 |
| *G. gallus* | GRCg7b | 1,053 | Ensembl 115 | ERR7015103, re-CCS | Sequel II Iso-Seq | 1,067,066 | 0.41% | 9,775 | galloanserae_odb12 |
| *D. rerio* | GRCz11 | 1,679 | Ensembl 115 | SRR32588721 | Sequel II Iso-Seq | 6,332,434 | 1.47% | 102,603 | cyprinodontiformes_odb12 |
| *M. musculus* C57BL/6J | GRCm39 | 2,728 | Ensembl 115 | ERR14205062 (brain) + ERR15371799 (kidney) | Sequel IIe Iso-Seq | 8,083,958 | 0.21% | 53,011 | eutheria_odb12 |
| *H. sapiens* WTC11 | GRCh38 | 3,292 | Ensembl 115 | 9 LRGASP FLNC runs | Sequel II FLNC | 7,629,668 | 0.63% | 44,765 | eutheria_odb10 |

### Evaluation of the pipeline design

The final pipeline design was established by evaluating five key decisions, each against a plausible alternative (Fig. 1). We assessed each alternative on the eight benchmark species before comparing SQANTI-evidence with other tools. When the alternatives offered complementary advantages, both were retained, with the default selected according to the benchmark results, allowing users to adapt the workflow to their needs.

### IsoQuant provides a more precise basis for annotation than IsoSeq3

In our previous work, transcriptome reconstruction relied on IsoSeq3 [21]. We therefore evaluated whether IsoQuant, which performed well without a reference annotation in LRGASP [13], could provide a more reliable basis for genome annotation. We compared both tools using the same reads and their unfiltered transcript outputs, thereby assessing reconstruction performance independently of subsequent curation (Fig. 2, Table S1).

**Figure 2.**
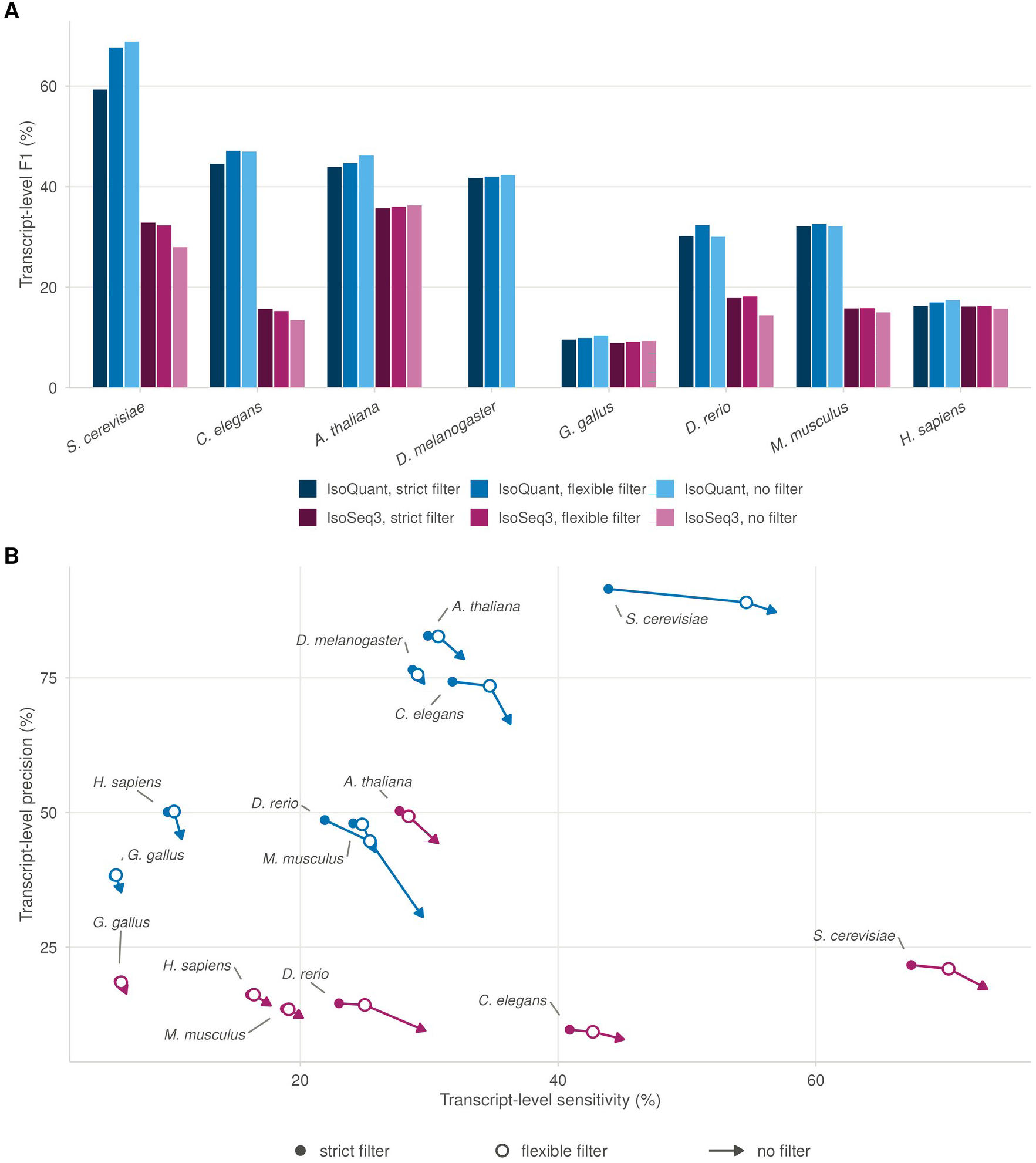
| Reconstruction method and curation filter, measured on the curated layer. gffcompare of Tier 1 alone on the CDS, transcript level (exact intron chain), against the CDS projection of the Ensembl reference; eight species with IsoQuant and seven with IsoSeq3 (no IsoSeq3 run for the Nanopore *D. melanogaster* dataset). Blue, IsoQuant; magenta, IsoSeq3; lightness encodes the filter (dark, strict filter; mid, flexible filter (default); light, no filter). **A**, Transcript-level F1 per species and run. **B**, Transcript-level sensitivity against precision. For each species and reconstruction method the filled point is the strict filter, the open circle the flexible filter and the arrowhead the unfiltered reconstruction, so each arrow traces the change from the strict to the flexible filter and then to no filter. IsoQuant is the more precise method in every species, and the flexible filter recovers sensitivity along the arrow at a precision cost below one point in six of eight species.

IsoSeq3 yielded 1.7 to 10.4 times as many coding transcript models as IsoQuant (194,378 against 37,286 in human; Table S1). However, IsoSeq3 transcript-level precision did not exceed 44.3% in any species, whereas IsoQuant reached 87.2% (Fig. 2B). The additional models increased transcript-level sensitivity in yeast, worm and human, by 6.9–16.4 percentage points, changed it by less than 0.5 points in chicken and zebrafish, and reduced it in Arabidopsis and mouse; nowhere did the gain compensate for the loss of precision. Consequently, IsoQuant achieved the higher transcript-level F1 in all seven species evaluated with both tools, by a margin that was smallest in chicken (1.1 percentage points) and largest in yeast (41 percentage points), the species in which IsoSeq3 had inflated the transcriptome most (Fig. 2A). At the locus level the two methods were within four percentage points of each other in five species (Fig. S1): they find largely the same genes and differ in the transcript structures they report. Because the annotation is delivered as transcript models, transcript-level accuracy is the relevant criterion, and IsoQuant is the default.

### A flexible curation filter for unannotated genomes

Because SQANTI-evidence is designed to be configurable, we evaluated several versions of the curation filter to determine how they affected the curated layer. We first applied a six-rule filter previously used to identify transcript artifacts from features independent of the reference annotation [21]. However, requiring all six conditions to be met proved too restrictive. Applied to the IsoQuant transcriptomes, the strict filter increased transcript-level precision but reduced sensitivity in all species. Because sensitivity was the lower of the two measures in every species, its loss outweighed the gain in precision. Consequently, transcript-level F1 ended below that of the unfiltered layer in seven of the eight species, with zebrafish as the only exception (Fig. 2, Table S1).

Most of this loss was attributable to a single rule. The intra-priming condition flags transcripts as potential artifacts when more than 59% of the genomic sequence downstream of their 3′ end is adenine. This condition alone accounted for the majority of the reference-exact isoforms lost in five of the eight species, and in every species most of the isoforms it removed were the only isoform of their gene, leaving no model for that gene in the curated layer (Fig. S2). Because the condition evaluates only the 3′ end, its ability to identify artifacts does not necessarily reflect the accuracy of the coding sequence on which annotation performance is assessed.

Read support helped distinguish genuine isoforms from artifacts among transcripts flagged by intra-priming. In five of the six animal datasets, with the Nanopore fly dataset as the exception, the proportion of flagged isoforms that matched a reference transcript exactly increased with the number of supporting full-length reads. In worm, this proportion rose from 4.5% in the lowest read-count group to 51.5% in the highest (Fig. S3A). We therefore evaluated alternative rule sets using the SQANTI3 classification tables, before gene prediction, to determine whether read support could mitigate the loss of genuine transcripts (Fig. S4; rejected alternatives are described in Supplementary Note S2). The selected alternative retained transcripts that failed exactly one of three conditions: intra-priming, PSAURON or canonical junctions; provided they were supported by at least ten full-length reads. This increased the proportion of reference-exact isoforms retained in every species, from a lower bound of 74% to at least 87.8%, while reducing precision by no more than 2.7 percentage points. We therefore adopted this flexible filter as the default in SQANTI-evidence.

In the curated layers, the flexible filter restored transcripts that the strict filter had removed in every species, most of all in zebrafish, where 4,956 were restored (Table S1; Fig. S3C). Sensitivity increased in all eight species, by up to 10.7 percentage points in yeast, whereas precision decreased by more than one percentage point only in yeast and zebrafish (Table S1). Consequently, transcript-level F1 increased in every species. Relative to the flexible filter, removing it altogether decreased precision by more than it increased sensitivity in every species except yeast. In zebrafish, precision decreased by 14.1 percentage points (Table S1).

Zebrafish, which had the noisiest of the eight transcriptomes, highlighted the limitations of a fixed read-support threshold. The flexible filter restored 891 reference-exact isoforms but also admitted 2,750 non-exact isoforms of annotated genes and 916 intergenic or intronic transcripts (Fig. S3C). As a result, transcript-level precision in the curated layer decreased from 48.6% to 44.7%. The optimal stringency therefore depends on transcriptome quality, which cannot be assessed directly against a reference annotation in the intended use case. To address this limitation, SQANTI-evidence reports a reference-free diagnostic before filtering: the fraction of reconstructed models classified as non-coding. This fraction was lowest in *Arabidopsis* (2.6%) and highest in zebrafish (60.8%), and it correlated with the reference-based fraction of spurious models across the eight species (Spearman’s ρ = 0.88; Fig. S3B).

Finally, we assessed whether curation could compensate for the lower precision of IsoSeq3-based reconstruction (Fig. 2). Applied to the IsoSeq3 transcriptomes, the flexible filter removed up to 44% of the models and raised transcript-level precision by no more than 5.0 percentage points. However, precision after filtering remained below 50% in every species, and the flexible and strict filters differed by less than 0.5 percentage points in transcript-level F1: the exceptions that recover genuine isoforms from IsoQuant have nothing to recover from IsoSeq3. Together, these results show that the filter determines the precision of the curated layer when the reconstruction is precise, but cannot overcome the limitations of an imprecise reconstruction.

### Hint types, rather than weights, determine the value of long-read evidence

The default AUGUSTUS hint weights were developed for transcriptomes reconstructed from short-read RNA-seq data, and their suitability for long-read evidence could not be assumed. We therefore evaluated both hint types and weights using human chromosome 19 and the entire *Arabidopsis* genome (Fig. S5, Table S2). Hint selection substantially affected prediction accuracy, whereas changes in weight had little effect.

Overall, long-read hints improved gene prediction. Relative to protein hints alone, the default hint set increased locus-level F1 by 6.5 percentage points in human and 1.5 percentage points in *Arabidopsis*. The smaller gain in *Arabidopsis* reflected its higher baseline accuracy with protein hints alone (locus-level F1, 67.5 versus 53.0 in human). However, the contribution depended on hint type. Intron and CDS hints, either individually or in combination, ranked highest among the eight combinations evaluated. All combinations containing exon hints performed worse than the default, and in human, intron-plus-exon and exon-only hints gave poorer results than the protein-only control. Adding CDS start and stop hints to the intron-plus-CDS combination did not change performance. These results indicate that hints describing features directly represented in the predicted gene models provide the most useful long-read evidence.

Bonus values had little effect on performance: varying the long-read bonus from 10⁴ to 10¹⁵ changed locus- and transcript-level F1 by no more than 0.3 percentage points. By contrast, the malus strongly affected prediction accuracy. Values below 1 penalize features lacking hint support, with the penalty accumulating across a gene’s exons. At a malus of 0.9, locus-level F1 decreased from 59.5 to 43.2 in human and from 69.0 to 31.9 in *Arabidopsis*, and at 0.5 the prediction quality scores collapsed in both genomes. Genes with no expression evidence were particularly affected because they lack long-read hints, despite being the primary targets of the predicted layer. We therefore adopted a default configuration using intron, CDS, and CDS start and stop hints, with a single weight file across species. All hint types and weights remain user-configurable.

### The predicted layer increases sensitivity at the cost of precision

We next assessed the contribution of the predicted layer to the final annotation by scoring the same runs on the curated layer alone and on the complete annotation (Fig. S6, Table S1). Starting from the curated layer generated with IsoQuant and the flexible filter, adding the predicted layer increased locus-level sensitivity in every species, by 7.4 percentage points in mouse and by up to 26.9 in fly. However, locus-level precision was reduced by 2.0 percentage points in yeast and by up to 34.2 in human. In the four compact genomes (yeast, worm, *Arabidopsis* and fly; 12–144 Mb), the loss in precision never exceeded the gain in sensitivity by more than 1.2 percentage points, whereas in the four vertebrate genomes (1.1–3.3 Gb) it was more than twice the gain (Table S1). The changes at transcript level followed the same pattern with smaller values. The predicted layer was also less precise than the curated layer. In the vertebrates, at most 15.6% of its transcripts were reference-exact, less than half the proportion in the curated layer, whereas in the compact genomes the proportion was at least 42.5% (Table S1). Thus, the precision of the complete annotation was largely determined by the predicted layer, and transcript reconstruction and filtering should be evaluated on the curated layer alone.

Unlike curated transcripts, multi-exon models in the predicted layer enter the final annotation without an evidence test. We therefore evaluated whether requiring at least one protein-supported intron per multi-exon predicted model would improve the annotation (Table 2). The requirement removed between 49% and 86% of the predicted models and raised the exact-CDS precision of the whole annotation in every species, most in chicken, by 15.8 percentage points. However, the removed models were not uniformly spurious. In *Arabidopsis*, worm and fly, more than 40% of them were reference-exact, compared with no more than 14% in the vertebrates. As a result, the requirement removed more than 10% of all reference-exact models in five of the seven species, up to 26.1% in chicken, and the loss remained below 5% only in mouse and human. Moreover, even after filtering, the exact-CDS precision of the whole annotation in the vertebrates did not exceed 55.3%. Because every model carries its tier attribute, the requirement can be applied after the run, whereas a model removed during the run cannot be recovered. We therefore deliver the predicted layer complete by default and leave this choice, which depends on the genome and on the intended use of the annotation, to the user (Supplementary Note S3).

**Table 2.** | Effect of an evidence requirement on Tier 2. Flexible-filter runs, seven species; every multi-exon Tier 2 model without a protein-supported intron is removed. Tier 2 models, Removed and Exact among removed count Tier 2 models. Exact before and Exact after are the exact-CDS precision of the whole annotation (Tier 1 plus Tier 2), the percentage of its models whose CDS intron chain matches a reference transcript, before and after the removal; Exact models lost is the percentage of all reference-exact models of the whole annotation that the requirement removes. The requirement is not applied by default; it can be applied after the run (Supplementary Note S3).

| **Species** | **Tier 2 models** | **Removed** | **Exact among removed** | **Exact before (%)** | **Exact after (%)** | **Exact models lost (%)** |
|---|---|---|---|---|---|---|
| *C. elegans* | 7,251 | 6,251 | 2,717 | 64.9 | 74.2 | 20.3 |
| *A. thaliana* | 10,376 | 6,758 | 2,725 | 75.4 | 88.2 | 14.3 |
| *D. melanogaster* | 6,919 | 4,771 | 2,162 | 72.7 | 85.0 | 19.3 |
| *G. gallus* | 11,631 | 9,677 | 1,208 | 25.9 | 41.7 | 26.1 |
| *D. rerio* | 25,200 | 15,317 | 1,654 | 30.5 | 39.5 | 11.1 |
| *M. musculus* | 10,677 | 5,256 | 722 | 49.8 | 55.3 | 3.6 |
| *H. sapiens* | 14,796 | 7,764 | 906 | 48.0 | 55.3 | 4.1 |

### Curated transcripts are retained as observed rather than re-predicted

We next evaluated how to integrate the curated and predicted layers into a single annotation. Curated transcripts can either be incorporated with their observed structures or supplied to AUGUSTUS as hints and re-predicted. To compare these alternatives, we matched each curated transcript of the eight benchmark species to the AUGUSTUS model predicted at the same genomic position before the layers were combined and determined whether their CDS intron chains agreed; UTRs were not compared because AUGUSTUS was run without UTR prediction (Fig. S7).

AUGUSTUS reproduced the coding structure of the curated transcript, or agreed that it was single-exon, in more than 96% of cases in yeast and *Arabidopsis* but only in 66.2% in human (Fig. S7). In the mammals, almost all of the disagreement consisted of AUGUSTUS models that shared some introns with the observed transcript but matched none of its isoforms, which is the expected outcome when the hints of several isoforms at one locus are combined into a single prediction. Structural mismatches and missing models accounted for no more than 3% of transcripts in any species. Re-prediction would also discard the UTRs, which nearly every curated transcript carries. Introns located only in the UTRs distinguished the observed transcript from the AUGUSTUS model in a quarter of the curated transcripts of *Arabidopsis* and in more than a third of those of chicken, mouse and human (Fig. S7).

When compared with the reference annotation at the same loci, curated transcripts matched the reference exactly more often than the AUGUSTUS models they replace in seven of the eight species (Table S3). In human, the curated layer also recovered 4,105 reference transcripts missed by AUGUSTUS, whereas AUGUSTUS recovered only 489 missed by the curated layer. Yeast was the exception, because AUGUSTUS already reproduced nearly all curated transcripts in this species. Together, these results support incorporating curated transcripts into the final annotation with their observed structures and UTRs intact, rather than re-predicting them.

We next assessed whether the rule used to combine the two layers, which discards every AUGUSTUS model that overlaps a curated gene on the same strand, removed valid gene structures, by comparing every discarded model with the reference annotation (Table S4). In every species, at least 69% of the discarded models had an intron chain already represented in the curated layer. Most of the remaining multi-exon models had incorrect structures: across the benchmark species, 1,716 matched reference transcripts absent from the curated layer, whereas 15,015 matched no reference transcript. Retaining these models would therefore have introduced 6–14 spurious models for every additional reference-supported isoform recovered. None of the 4,926 single-exon models discarded in the six animal genomes was reference-exact, whereas 72% were in yeast and 38% in *Arabidopsis*, in which single-exon genes are prevalent. Thus, the combination rule removed few correct structures other than the single-exon models of these two species.

Together, these evaluations established the default configuration of SQANTI-evidence: transcript reconstruction with IsoQuant, curation with the flexible filter, AUGUSTUS prediction guided by intron, CDS, and CDS start and stop hints using a single weight file across species, and direct incorporation of curated transcripts with their observed structures preserved. AUGUSTUS predictions are retained only at loci without a curated gene.

### Comparison with BRAKER3 and BRAKER4

#### Overall accuracy and completeness

We compared the final SQANTI-evidence annotations with those generated by BRAKER3 and BRAKER4 in the seven species for which both BRAKER versions could be run (Fig. 3; Tables S1 and S5). BRAKER4, run with OrthoDB, achieved the highest locus-level sensitivity in all seven species. Its advantage over SQANTI-evidence was smallest in zebrafish (14.1 percentage points) and largest in chicken (24.4). It was also more precise than SQANTI-evidence in every species, by more than 10 percentage points (Fig. 3). BRAKER3, which used only information derived from the same long-read data, showed no consistent difference in sensitivity from SQANTI-evidence. It was more sensitive in four species and less sensitive in three, never by more than 7.1 percentage points. However, it was more precise than SQANTI-evidence in every species, by more than 13 percentage points (Fig. 3).

**Figure 3.**
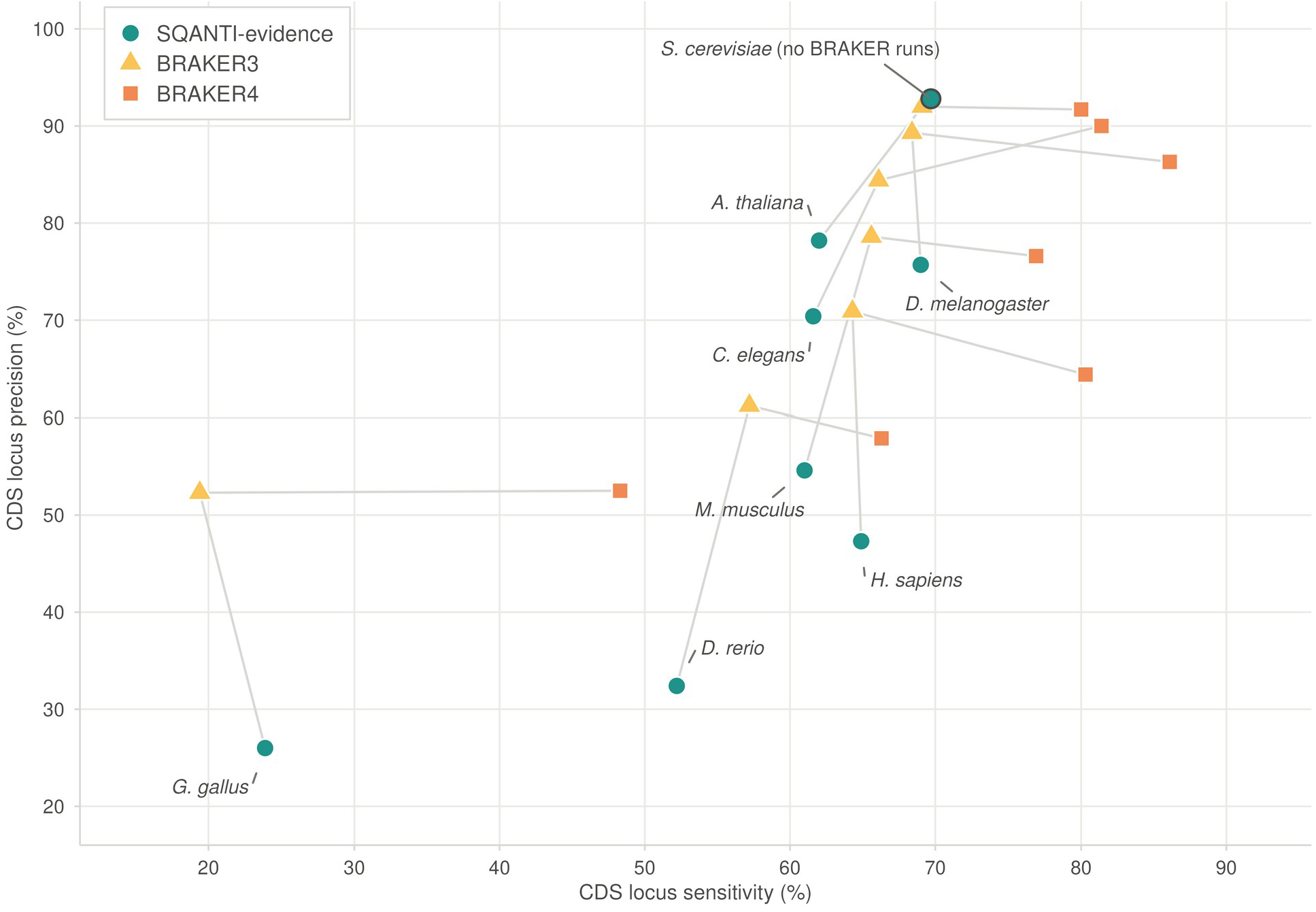
| Aggregate accuracy of the three annotations. CDS locus sensitivity against CDS locus precision for SQANTI-evidence with the flexible filter, BRAKER3 run on the same FLNC reads and on the proteins of the unfiltered transcriptome, and BRAKER4 run with the OrthoDB v12 clade proteins, eight species; *S. cerevisiae* has no BRAKER point because GeneMark-ETP fails on it. Gray segments join the three annotations of one species. BRAKER4 lies to the upper right of SQANTI-evidence in every species. BRAKER3, given the same reads and sample-derived proteins, is comparable in sensitivity and higher in precision.

All three methods performed poorly in chicken, probably because the dataset represented a single tissue and developmental stage. The reads covered only approximately one-third of the reference coding genes (5,274 of 17,007), limiting the available transcriptomic evidence.

The GAQET2 metrics yielded a different ranking of annotation completeness (Fig. 4A; Fig. S8). SQANTI-evidence achieved higher BUSCO completeness than BRAKER3 in five of the seven species, equal in mouse and lower in worm, and its margin was largest in chicken, at 29.6 percentage points. It also left fewer genes missing according to OMArk in all seven species, again with the largest margin in chicken (41.9 percentage points). BRAKER4 was the most complete annotation on both measures in every species. Thus, of the two annotations generated from the same evidence, SQANTI-evidence produced a more complete core proteome than BRAKER3.

**Figure 4.**
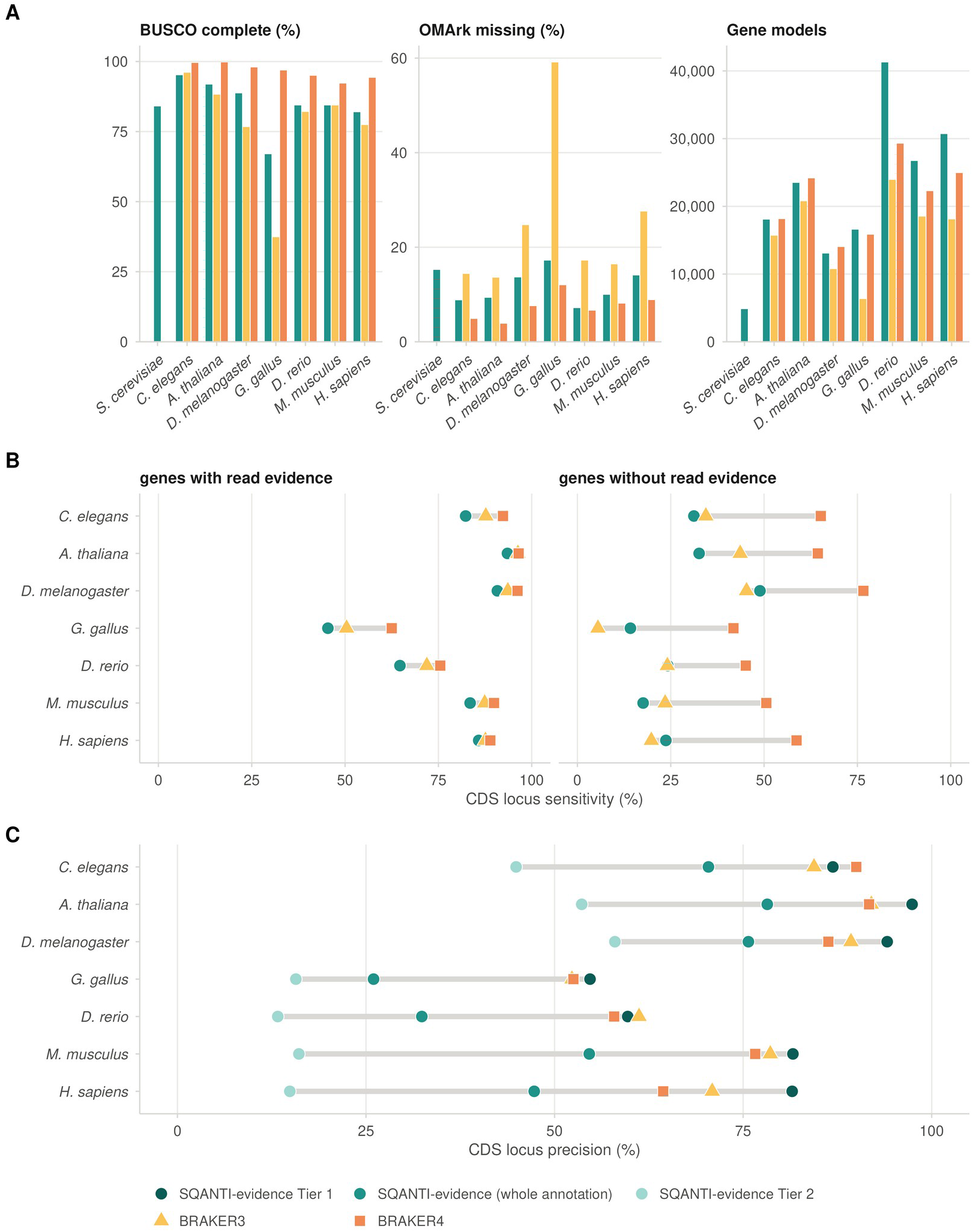
| Where the annotations differ: proteome completeness, sensitivity by read evidence, precision by source. **A**, GAQET2 metrics of the whole annotations: annotation BUSCO completeness, OMArk missing genes and number of gene models. Panel A also includes *S. cerevisiae*, for which no BRAKER annotation could be produced. **B**, CDS locus sensitivity at reference coding genes overlapped by at least one IsoQuant model before filtering and at genes overlapped by none. A gene is considered to be recovered when one predicted transcript reproduces its CDS intron chain. The three annotations are closer at genes with read evidence; the gap to BRAKER4 lies at genes without it, where BRAKER3 with proteins from the same long reads has no consistent advantage. **C**, CDS locus precision of the models grouped by source: SQANTI-evidence Tier 1, whole annotation, Tier 2, BRAKER3 and BRAKER4. Tier 1 is the most precise set in five of seven species and Tier 2 the least in all seven. SQANTI-evidence was run with the flexible filter, and BRAKER3 with the same reads, throughout.

This increased completeness was accompanied by greater redundancy. SQANTI-evidence contained more duplicated BUSCOs than BRAKER3 in every species, with the largest difference in human, 15.7% compared with 4.8% (Fig. S8). In zebrafish, mouse and human, it also had a lower proportion of OMArk-consistent proteins; in zebrafish, for example, this fraction was 83.8% compared with 94.9% for BRAKER3. SQANTI-evidence additionally predicted more transcripts per gene (Fig. S9). In human, the complete annotation averaged 1.51 transcripts per gene, compared with 1.32 and 1.44 for BRAKER3 and BRAKER4, respectively, while the curated layer alone averaged 1.88.

### Genes with and without read evidence

To identify the source of the sensitivity differences, we stratified the reference coding genes according to whether they overlapped at least one IsoQuant model before filtering, hereafter genes with and without read evidence (Fig. 4B, Table S6). At genes with read evidence, both BRAKER versions were more sensitive than SQANTI-evidence in every species: BRAKER4 exceeded SQANTI-evidence by no more than 17.1 percentage points, in chicken, and the advantage of BRAKER3 was smaller still in every species (Table S6).

At genes without read evidence, the two BRAKER versions differed. BRAKER4 was more sensitive than SQANTI-evidence in every species, by more than 20 percentage points in each (Table S6). Weighted by the number of genes in each stratum (Table S6), these genes accounted for most of the overall sensitivity advantage of BRAKER4 in every species except zebrafish, where the genes with read evidence contributed more. BRAKER3, which relied on protein evidence derived from the same long-read data, showed no consistent advantage over SQANTI-evidence at these genes. SQANTI-evidence recovered more of them in fly, chicken and human, fewer in worm, *Arabidopsis* and mouse, and nearly the same proportion in zebrafish, and no difference exceeded 11.0 percentage points. Therefore, when both methods relied on the same evidence, neither recovered consistently more of the genes without read evidence, and the advantage of BRAKER4 at these genes is largely attributable to the external protein evidence it obtains from OrthoDB.

### Precision of the curated and predicted layers

To determine the contributions of each layer to annotation precision, we evaluated the curated and predicted models separately against the two BRAKER annotations (Fig. S10). At the locus level, the curated layer was the most precise gene set in five of the seven species (Fig. 4C; Table S5). Its locus-level precision exceeded that of BRAKER4 in every species except worm and that of BRAKER3 in every species except zebrafish. The comparison was less favorable when precision was assessed at the model level using exact-CDS matches. The curated layer outperformed BRAKER4 in five species, including human (63.8% versus 54.1%), whereas BRAKER3 achieved higher exact-CDS precision in every species except *Arabidopsis* and fly, by no more than 6.4 percentage points (Fig. S10). However, under the less stringent criterion of overlapping an annotated coding gene, the curated layer outperformed both BRAKER versions in all seven species (Table S5). This proportion was lowest in zebrafish, at 78.7%. Thus, even there most of the models that the exact-CDS criterion counted as false positives overlapped a coding gene with a different structure. The principal limitation of the curated layer was coverage: it recovered no more than 54.4% of all coding genes in any species, and only 13.2% in chicken, a lower proportion than either BRAKER version in every species (Table S5).

The predicted layer was the least precise of the four gene sets in every species (Fig. S10). To investigate the origin of this lower precision, we compared SQANTI-evidence predictions with the AUGUSTUS-derived models retained by TSEBRA [22] in each BRAKER annotation, stratifying models according to whether their introns had protein support (Fig. S11). Among models without protein support, the AUGUSTUS models retained by TSEBRA matched the reference exactly more often than the SQANTI-evidence predictions, by more than 28 percentage points in every species and in both BRAKER annotations (Fig. S11). This difference reflected the distinct model-selection strategies: BRAKER uses evidence support to select AUGUSTUS predictions, whereas SQANTI-evidence retains every multi-exon AUGUSTUS model outside the curated layer and applies an evidence filter only to single-exon predictions.

The genomic context of the non-exact models in the predicted layer also differed between species (Table S7). In worm, *Arabidopsis*, fly and chicken, at least 77% of the non-exact models overlapped a coding gene with a different structure. In mouse and human, models overlapping non-coding genes or pseudogenes were more frequent than models overlapping coding genes with a different structure, 43% versus 37% in mouse, and no more than 12.3% of the non-exact models of either species lay predominantly in soft-masked sequence. By contrast, in zebrafish the largest class of non-exact models lay predominantly in soft-masked sequence, at 39%, and models overlapping coding genes with a different structure accounted for 36%. These results indicate that a permissive predicted layer remains susceptible to spurious gene models in repeat-rich regions even when the genome has been soft-masked.

### Selection of annotation subsets by layer

Because every model carries a tier attribute, SQANTI-evidence allows users to select different annotation subsets after the run rather than providing a single fixed annotation. Figure 5 illustrates how locus-level sensitivity and precision changed in four species as the predicted layer was progressively restricted, from the complete default annotation (A) to the curated layer alone (D), followed by a subset that excluded curated loci overlapping no annotated gene from the precision denominator (E). Subsets A and D are selected directly with the tier attribute, B and C additionally use the protein-hint file written by every run (Supplementary Note S3), and E requires the reference annotation.

**Figure 5.**
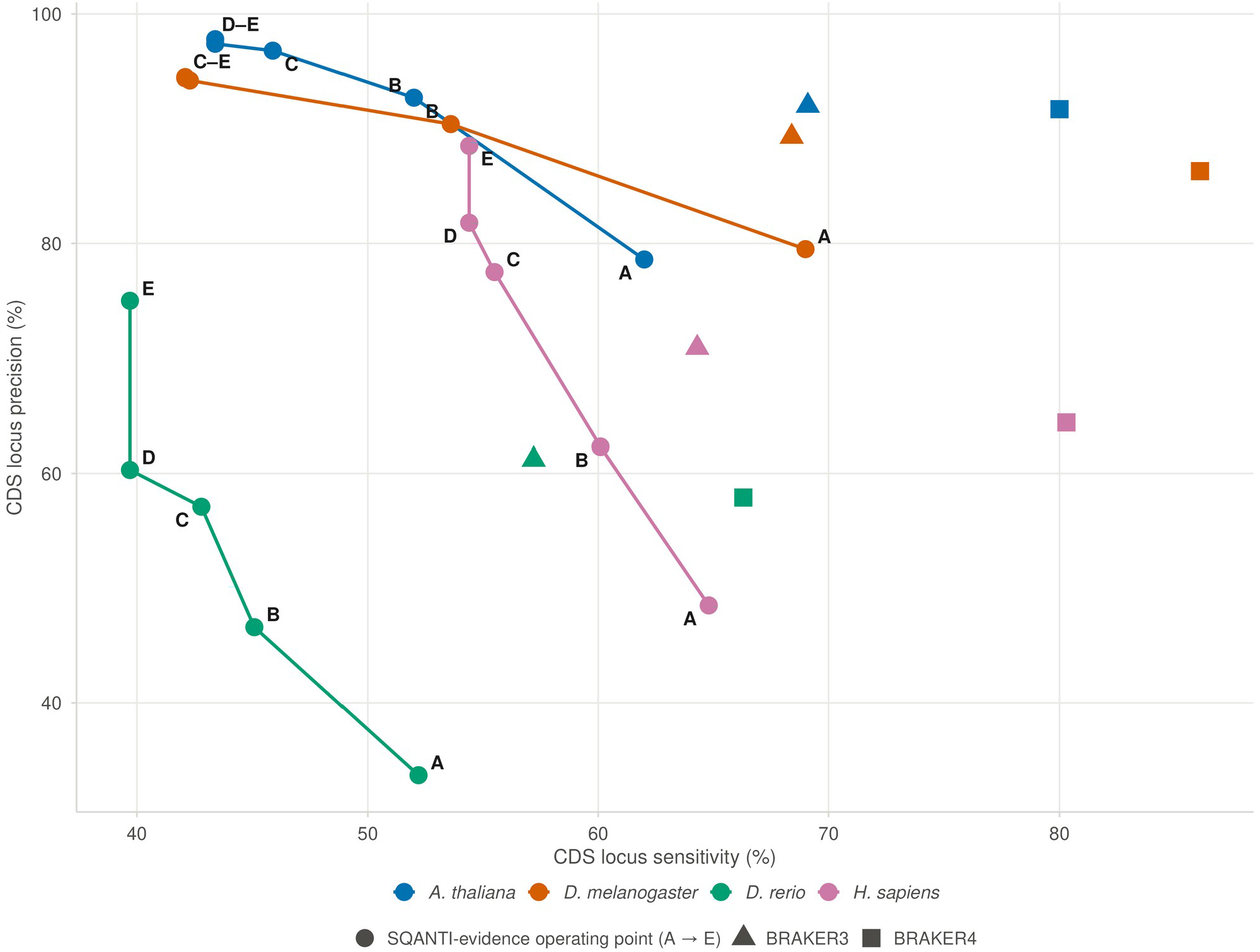
| Subsets of the SQANTI-evidence annotation. CDS locus sensitivity against precision as Tier 2 is restricted step by step, for *A. thaliana*, *D. melanogaster*, *D. rerio* and *H. sapiens*: A, default annotation (all Tier 2); B, Tier 2 restricted to models with every intron protein-supported; C, B without single-exon Tier 2 models; D, Tier 1 alone; E, D with the Tier 1 loci that overlap no annotated gene excluded from the precision denominator. Triangles and squares, BRAKER3 and BRAKER4 for the same species. Steps B–D are filters applied after the run (B and C use the protein hints written with every run); E requires the reference annotation.

In human, locus-level sensitivity and precision changed from 64.8% and 48.5% in the complete annotation to 54.4% and 81.8% in the curated layer, respectively. Excluding curated loci that overlapped no annotated gene increased precision further to 88.5%, without changing sensitivity (Table S5). This final subset distinguished two sources of apparent imprecision in the curated layer. Loci excluded from the denominator fell in unannotated regions, where the reference could not establish whether a model represented a novel gene or an artifact. The remaining unmatched loci overlapped annotated genes but had a different CDS structure, or corresponded to non-coding genes.

The BRAKER annotations occupied single points in this sensitivity–precision space because TSEBRA had already selected their gene models. The curated layer alone achieved higher precision than both BRAKER annotations in five of seven species, with worm and zebrafish as exceptions, whereas the default SQANTI-evidence annotation provided the highest sensitivity among its own configurations (Table S5). Users can move between these trade-offs simply by filtering on the tier attribute. Figure S12 presents the corresponding subsets at transcript level, using exact-CDS precision.

### Isoform recovery

Isoforms represent different types of evidence in SQANTI-evidence and BRAKER. In the curated layer of SQANTI-evidence, each isoform corresponds to a transcript sequenced end to end, passing quality control and incorporated into the annotation without structural modification. In BRAKER and the predicted layer of SQANTI-evidence, isoforms are reconstructed from the combined hints derived from reads at each locus and may therefore differ from the observed transcripts (Fig. S7). This distinction was particularly evident in genes with extensive alternative splicing (Fig. S13). Among reference genes with multiple annotated CDS isoforms, SQANTI-evidence recovered at least two isoforms exactly in approximately twice as many genes as either BRAKER version in mammals. In human, this proportion was 29.8%, compared with 14.6% for BRAKER3 and 16.0% for BRAKER4; in mouse, it was 23.4%, compared with 11.9% and 12.6%, respectively. Differences were smaller in the other species. SQANTI-evidence exceeded both BRAKER versions by less than 5 percentage points in fly, *Arabidopsis* and zebrafish, whereas all three methods recovered exact isoforms in fewer than 5% of chicken genes with multiple annotated CDS isoforms. Worm was the exception, with SQANTI-evidence recovering multiple exact isoforms in 12.5% of genes, compared with 17.2% for BRAKER3 and 21.6% for BRAKER4. This difference may have reflected the compact, operon-rich transcriptome of *C. elegans*, in which resolving alternative isoforms may offer less benefit. BRAKER4 recovered more genes with at least one exact isoform in every species, consistent with its higher overall gene recovery. However, only SQANTI-evidence explicitly distinguishes isoforms directly observed in the sample from those inferred computationally.

### Read-derived UTRs place 3′ ends close to the poly(A) site

Because curated transcripts retain the original read boundaries, which may extend beyond the CDS, SQANTI-evidence preserves read-derived UTRs wherever transcript evidence is available. Accordingly, nearly every curated transcript carried both UTRs, 97.3% or more in every species. The predicted layer, by contrast, contains only CDSs, because AUGUSTUS was not run in UTR-prediction mode given the absence of UTRs from its training set. We therefore compared SQANTI-evidence UTRs with the experimentally informed UTR annotations generated by BRAKER4 (Fig. 6 and Fig. S14), assessing the 3′ and 5′ ends separately.

**Figure 6.**
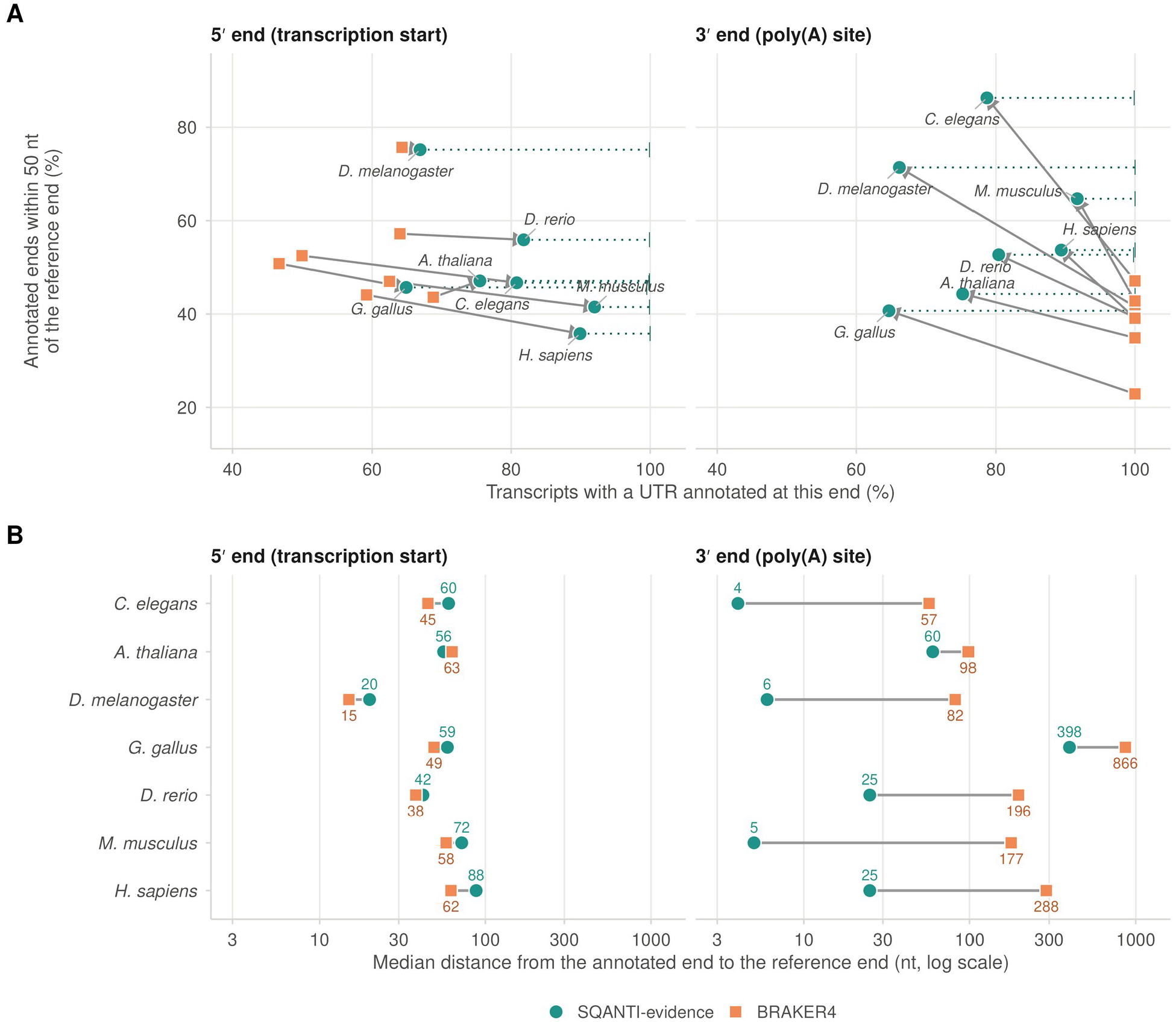
| Read-derived UTRs place the poly(A) site more accurately than an assembly of the same reads. UTR ends of the whole SQANTI-evidence annotation and of BRAKER4’s experimental *stringtie2utr* output, scored on the reference transcripts whose CDS both tools reproduce exactly, at the ends where the reference annotates a UTR, seven species. **A**, Share of these transcripts with a UTR annotated at the end (coverage) against the share of annotated ends within 50 nt of the reference end (accuracy). Each arrow runs from BRAKER4 to SQANTI-evidence for one species; the dotted extension marks the coverage of Tier 1 alone, 99.5–100% because Tier 2 is CDS-only. In the eight benchmark species, 97.3–99.7% of curated transcripts carry both UTRs. **B**, Median distance between annotated and reference ends, log scale, values at the points. At the 3′ end SQANTI-evidence is closer in all seven species, eight-to thirty-five-fold in five of them; at the 5′ end it annotates more ends and BRAKER4 places them closer in most species, by a small difference.

At the 3′ end, SQANTI-evidence was more accurate than BRAKER4 in all seven species. In five of them, the median distance between SQANTI-evidence transcript ends and the annotated poly(A) site was 25 nt or less, an eight-to 35-fold reduction relative to BRAKER4 (Fig. 6B). Worm provided an informative independent validation: the median SQANTI-evidence distance was just 4 nt relative to the independently curated WormBase annotation [23], supporting the accuracy of the read-derived ends rather than merely their consistency with the pipeline itself. The proportion of 3′ ends within 50 nt of the site was higher for SQANTI-evidence in every species, between 41% and 86%, whereas BRAKER4 never exceeded 47% (Fig. 6A). BRAKER4 annotated a 3′ UTR for every transcript in this comparison, whereas SQANTI-evidence did so for 65% to 92% of them, depending on the species. However, BRAKER4 assigned UTRs irrespective of the evidence supporting the transcript end, and between half and four-fifths of its predicted 3′ ends fell short of the annotated poly(A) site (Fig. S14).

The pattern was reversed at the 5′ end. SQANTI-evidence annotated a 5′ UTR for at least 65% of transcripts in every species, whereas BRAKER4 never exceeded 69%, and within the curated layer nearly every transcript carried one (Fig. 6A). Nevertheless, BRAKER4 placed its 5′ ends closer to the reference annotation in six species, by a median difference of no more than 26 nt; in human, for example, the median distance was 62 nt for BRAKER4 and 88 nt for SQANTI-evidence (Fig. 6B). *Arabidopsis* was the exception. The signed distances indicated that this discrepancy was not attributable to transcript truncation, which would be expected to place read boundaries inside the annotated transcript [14]. Instead, SQANTI-evidence 5′ ends extended beyond the annotated start in two-thirds or more of the transcripts in worm, fly, zebrafish, mouse and human, with median offsets of +13 to +64 nt (Fig. S14). In chicken, ends occurred on both sides of the annotated start, whereas in *Arabidopsis* they were located predominantly inside it, and in that species both methods showed a mode approximately 100 nt downstream of the annotated 5′ end. BRAKER4 ends, by contrast, fell on either side of the annotated start with no consistent direction, extending beyond it in as few as 22% of transcripts in one species and as many as 81% in another.

Together, these results show that BRAKER4 recovered more genes, largely through its additional protein evidence source, whereas the curated layer of SQANTI-evidence provided the highest locus-level precision in five of seven species, recovered more exact isoforms in mammals and placed 3′ transcript ends closer to the annotated poly(A) site in every species.

### Annotation of ERGA species

Finally, we applied SQANTI-evidence to 13 datasets from the ERGA initiative (Table 3, Fig. 7). Across the six ERGA pilot genomes, the curated layer contained from 7,025 genes in *Xylocopa* to 33,504 in *Salvelinus*. BUSCO completeness of the resulting annotations ranged from 78.2% in *Bufotes*, assessed against a substitute lineage, to 92.3% in *Zingel*.

**Figure 7.**
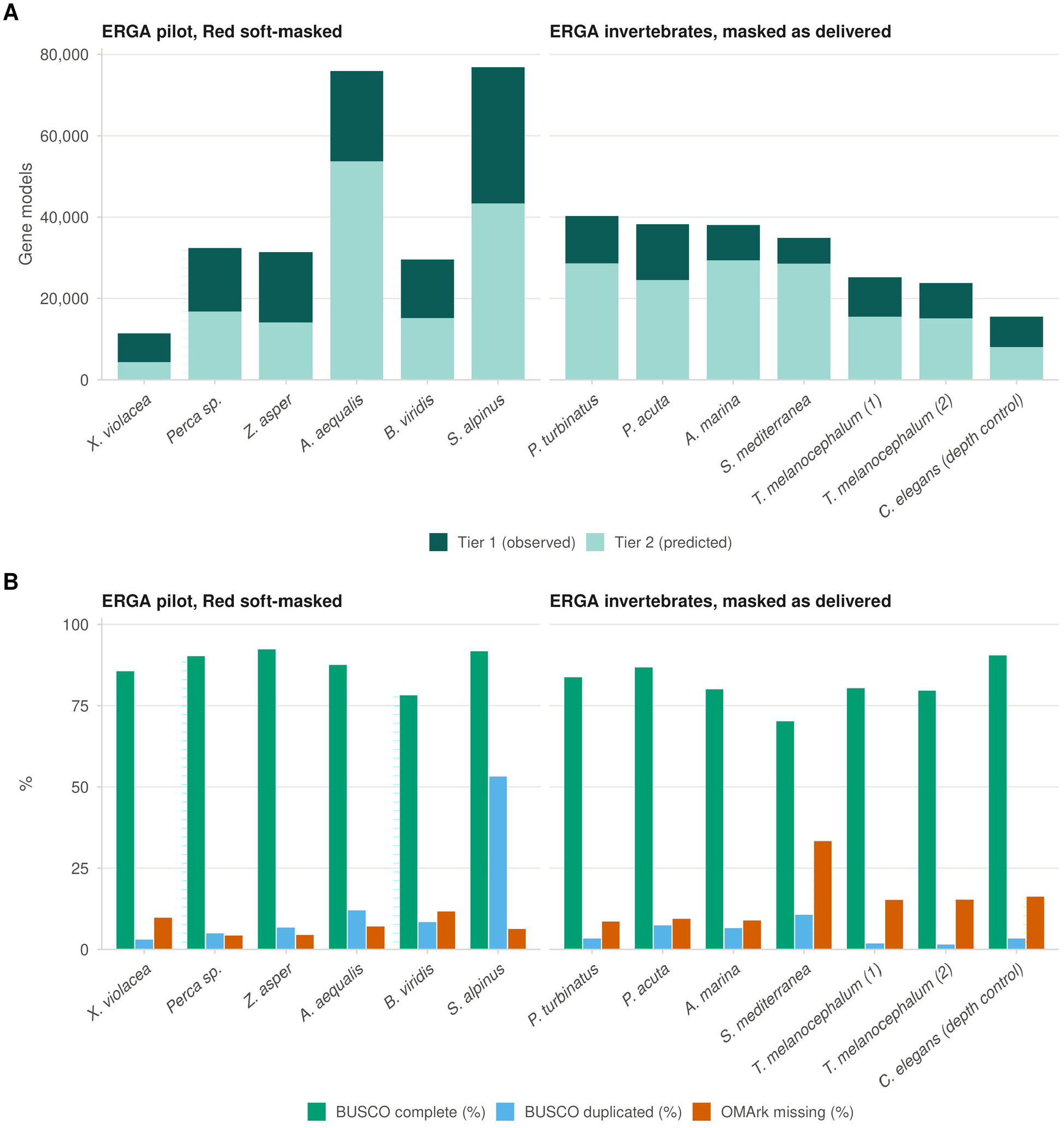
| Thirteen ERGA datasets annotated with SQANTI-evidence. Left group, the six ERGA pilot genomes after Red soft-masking; right group, seven invertebrate datasets on genomes delivered soft-masked, including a *C. elegans* library sequenced 34-fold shallower than the benchmark worm as a depth control. **A**, Gene models per tier: Tier 1, observed; Tier 2, predicted (pale teal). **B**, Annotation BUSCO completeness and duplication and OMArk missing genes, in percent. *B. viridis* is scored on tetrapoda_odb12 for lack of an amphibian lineage; the 53.2% duplicated BUSCOs of *S. alpinus* are consistent with the salmonid whole-genome duplication.

**Table 3.** | ERGA annotations. The deposited annotations: the six pilot genomes after Red soft-masking (first six rows) and the seven invertebrate datasets on the genomes as delivered. Tier 1 transcripts, curated transcripts of the annotation. BUSCO C, annotation BUSCO completeness on the lineage given in the table; OMArk missing, fraction of the clade’s HOGs absent from the annotated proteome. The last row is the *C. elegans* depth control, a shallow ERGA library annotated on the same assembly as the benchmark *C. elegans* dataset.

| **Species** | **Genome (Mb)** | **Soft-masked (%)** | **FLNC reads** | **Tier 1 genes** | **Tier 1 transcripts** | **Tier 2 genes** | **Total genes** | **BUSCO lineage (odb12)** | **BUSCO C (%)** | **OMArk missing (%)** |
|---|---|---|---|---|---|---|---|---|---|---|
| *Xylocopa violacea* | 1,021 | 80.6 | 3,471,737 | 7,025 | 13,894 | 4,391 | 11,416 | apoidea | 85.6 | 9.7 |
| *Perca sp. ‘yellow fin Alpine’* | 1,006 | 42.4 | 3,896,845 | 15,604 | 20,964 | 16,772 | 32,376 | actinopterygii | 90.2 | 4.3 |
| *Zingel asper* | 964 | 43.3 | 2,414,109 | 17,269 | 24,452 | 14,109 | 31,378 | actinopterygii | 92.3 | 4.4 |
| *Alopecurus aequalis* | 2,849 | 76.8 | 7,031,733 | 22,246 | 29,409 | 53,694 | 75,940 | poales | 87.5 | 7.0 |
| *Bufotes viridis* | 3,880 | 65.3 | 3,466,921 | 14,354 | 21,796 | 15,216 | 29,570 | tetrapoda | 78.2 | 11.7 |
| *Salvelinus alpinus* | 2,997 | 67.3 | 7,273,335 | 33,504 | 55,489 | 43,381 | 76,885 | actinopterygii | 91.7 | 6.3 |
| *Phorcus turbinatus* | 952 | 30.9 | 1,122,003 | 11,621 | 13,100 | 28,665 | 40,286 | mollusca | 83.7 | 8.5 |
| *Physella acuta* | 702 | 42.2 | 2,185,518 | 13,695 | 19,434 | 24,547 | 38,242 | mollusca | 86.8 | 9.4 |
| *Arenicola marina* | 869 | 33.8 | 1,375,632 | 8,620 | 10,661 | 29,414 | 38,034 | lophotrochozoa | 80.0 | 8.9 |
| *Schmidtea mediterranea* | 854 | 55.7 | 793,918 | 6,357 | 6,663 | 28,543 | 34,900 | lophotrochozoa | 70.2 | 33.3 |
| *Tetrastemma melanocephalum*, library 1 | 522 | 34.9 | 1,463,589 | 9,673 | 13,446 | 15,505 | 25,178 | lophotrochozoa | 80.4 | 15.2 |
| *Tetrastemma melanocephalum*, library 2 (Revio) | 522 | 34.9 | 960,516 | 8,663 | 12,212 | 15,116 | 23,779 | lophotrochozoa | 79.6 | 15.3 |
| *Caenorhabditis elegans* | 100 | 35.7 | 531,656 | 7,435 | 8,040 | 8,094 | 15,529 | chromadorea | 90.5 | 16.2 |

None of the six pilot genomes had been repeat-masked. We therefore annotated each assembly both before and after soft-masking with Red (Table S8, Fig. S15). The two annotation layers responded markedly differently to masking. Soft-masking, which required a few hours per genome, removed more than half of the predicted genes in every genome, up to 88%, and most of the single-exon transcripts. In contrast, the curated layer changed by no more than four genes, the number of transcripts containing both UTRs remained unchanged, and BUSCO completeness remained stable or increased in five of the six genomes, with a decrease of just 0.1 percentage points in the remaining genome. Thus, with the same reads and annotation workflow, soft-masking eliminated most predicted gene models without reducing BUSCO completeness by more than 0.1 percentage points. These findings indicate that unmasked repeats account for a substantial proportion of excess predictions in newly assembled genomes. We therefore recommend soft-masking assemblies before annotation and deposited the annotations generated from the masked genomes.

*Salvelinus* was an exception to the general pattern of duplication, with 53.2% duplicated BUSCOs after masking (Fig. 7B), consistent with the whole-genome duplication shared by salmonids [24]. The seven ERGA invertebrate datasets [25] had already been soft-masked and contained fewer reads than any of the pilot datasets (Table 3). In the six datasets other than the *C. elegans* depth control, the predicted layers were correspondingly larger, containing 1.6–4.5 times as many models as the curated layers. Annotation BUSCO completeness of these six datasets ranged from 79.6% to 86.8%, except in the planarian *Schmidtea mediterranea*, which reached 70.2% completeness and had 33% of OMArk genes missing. This species was the most divergent of the six and had the fewest reads.

The *C. elegans* dataset provided a direct assessment of sequencing depth. It used the same species and assembly as the benchmark worm dataset but contained 531,656 reads, compared with 18.1 million in the benchmark, and its curated layer contained 8,040 transcripts, compared with 13,376 transcripts in the higher-coverage benchmark dataset. Across the complete annotation, CDS-level locus sensitivity and precision were 47.1% and 62.0%, respectively, compared with 61.6% and 70.4% in the benchmark. For the curated layer alone, sensitivity and precision were 30.9% and 87.4%, compared with 46.3% and 86.9%. These results show that reduced sequencing depth primarily compromised the sensitivity of the curated layer, while its precision remained largely unchanged.

## Discussion

The rapid growth in genome assemblies generated by biodiversity initiatives has shifted the bottleneck from genome assembly to annotation. Long-read RNA sequencing captures full-length transcripts, yet current annotation pipelines use these data primarily as hints for gene prediction. In this study, we present SQANTI-evidence, a workflow that extends our previous strategy [21] into a reference-free annotation pipeline. SQANTI-evidence incorporates sequenced transcripts, including their isoforms and UTRs, into the annotation, labels them to distinguish directly observed transcripts from predicted models, and uses the same evidence to guide AUGUSTUS at loci not covered by the reads. Our results demonstrate that long-read transcriptomes can be curated without a reference annotation, providing a curated layer of precise gene models, isoforms and transcript ends that conventional evidence-driven predictors do not explicitly identify as directly observed.

Evaluation of the design choices showed that the original rule set was too restrictive for annotation, largely because the intra-priming condition evaluates a transcript’s 3′ end, whereas accuracy was assessed at the CDS level. Read support helped distinguish genuine transcripts from artifacts among the flagged models, allowing the flexible filter to recover most of the lost sensitivity at a small cost in precision. However, curation could not compensate for inaccurate transcript reconstruction, which is why IsoQuant is the default. IsoSeq3 remains available as an alternative because reference annotations are incomplete, and the benchmark cannot distinguish its artifacts from genuine transcripts that have not yet been annotated. Among the hint types tested, exon hints reduced prediction accuracy because, with UTR prediction disabled, they define transcript boundaries that include UTR sequence that a CDS-only model cannot reproduce. Re-predicting curated transcripts would also discard information established directly by the reads. Consequently, an isoform in a SQANTI-evidence annotation represents a different type of evidence from an isoform in a BRAKER annotation.

BRAKER4 with OrthoDB was more accurate overall in every genome in which it could be run, with much of its advantage arising at genes not covered by the reads. When supplied with proteins derived from the same long reads, BRAKER3 showed no consistent advantage at these loci, indicating that BRAKER4’s improvement largely derives from its external protein database. SQANTI-evidence deliberately relies only on proteins from the sample, allowing it to run on any genome for which transcript reads are available. Its main precision deficit lies in the predicted layer. TSEBRA removes AUGUSTUS models with insufficient evidence support, a strategy BRAKER can use because GeneMark-ETP provides a second source of predictions. SQANTI-evidence has only one predictor, making this selection more consequential. In a reference-free workflow, the only evidence available at loci not covered by reads comes from aligning proteins derived from the sample’s own transcripts. By construction, these proteins support paralogs of expressed genes but provide no direct support for genes that are not expressed. Requiring such support would therefore remove correct *ab initio* models at precisely the unexpressed genes that the predicted layer is intended to recover, as our tests demonstrated. Future versions could address this limitation by validating predictions with external proteins where available and by developing genome-dependent curation of the predicted layer, potentially using the noise diagnostic as a starting point.

The performance of all tools is constrained by the completeness of the reference annotations, which are still being updated to reflect the isoform diversity revealed by long reads [12]. Transcripts that match no annotated structure are counted as errors, whether they are artifacts or genuine, previously unannotated isoforms. Nevertheless, most curated loci overlapped an annotated coding gene (Table S5), and the read-supported loci in unannotated regions may include genes absent from the reference (Fig. 5). UTRs illustrate the value of directly observed transcripts. IsoQuant [10] places the 3′ end, when supported by a poly(A) tail, at the most frequent read end, positioning the curated model near the poly(A) site. It places the 5′ end at the furthest upstream supported read start, meaning that curated 5′ ends tend to extend beyond the annotated start rather than fall short of it. Because transcription start and termination sites account for many isoform differences between human tissues [26], and current predictors do not yet model these boundaries reliably, UTRs should be considered reliable only where supported by long reads, as in the curated layer.

The ERGA genomes, which had long-read data but no reference annotation, provided a direct test of the workflow in its intended setting. Soft-masking removed most predicted genes while leaving the curated layer largely unchanged. This difference reflects the distinct sources of the two layers: the curated layer is determined by the reads, whereas the predicted layer depends on the genome sequence supplied to the predictor. The types of errors in the predicted layer also varied between genomes. Among incorrect predicted models that did not overlap coding genes, human models were concentrated in pseudogenes, non-coding genes and unmasked intergenic regions, whereas in zebrafish many fell within sequence that was already soft-masked. Thus, a permissive prediction layer can also generate errors in repeat-rich regions of masked genomes. Other features of the deposited annotations reflect biology or data availability rather than annotation errors. The high duplication in *Salvelinus* is consistent with the salmonid whole-genome duplication, whereas the large predicted layers in invertebrates reflected their lower sequencing depth. The *C. elegans* depth-control experiment showed that lower depth reduces the sensitivity of the curated layer rather than its precision. The planarian *S. mediterranea*, the most divergent species and, apart from the depth control, the one with the fewest reads, illustrates the limits of annotation from a single, low-depth library.

This study has several limitations. Only one dataset was sequenced using Nanopore, the chicken dataset represented a single tissue, hint types and weights were evaluated on only one human chromosome and one small genome, and UTR predictions were compared with an experimental BRAKER output. Based on our results, we recommend soft-masking genome assemblies before annotation, checking read-alignment error rates, using IsoQuant with the flexible filter, and consulting the noise diagnostic when choosing filtering stringency. For libraries with high levels of noise, as in zebrafish, increasing the read-support threshold or using the strict filter is preferable when precision is more important than sensitivity. The curated layer is the appropriate choice when precision is the priority, whereas the full annotation is better suited to applications that require completeness. Where a protein database is available for the relevant clade, BRAKER4 can recover more genes; SQANTI-evidence, however, explicitly identifies the transcripts observed in the sample.

## Conclusions

SQANTI-evidence demonstrates that long-read transcriptomes can serve as a primary component of genome annotation rather than merely as evidence for gene prediction. As long-read RNA sequencing becomes increasingly common in biodiversity initiatives such as ERGA and the Earth BioGenome Project, directly observed transcripts could provide the foundation of new genome annotations, with gene prediction extending coverage to loci not reached by the reads. By distinguishing these two sources of information, SQANTI-evidence makes the origin of each gene model explicit and allows users to select annotation subsets according to their priorities for precision, completeness and transcript-level evidence.

## Methods

### The SQANTI-evidence workflow

SQANTI-evidence is implemented as a Snakemake workflow [27], with a Python wrapper script that controls the main options of the pipeline. It is designed to run either on a single-node workstation or in an HPC environment, where independent jobs are dispatched in parallel through the cluster scheduler. The workflow is end-to-end: the user supplies the long reads (PacBio full-length non-concatemer (FLNC) reads in BAM or FASTQ format, or Oxford Nanopore Technologies (ONT) reads) and the genome assembly, and the pipeline carries out transcriptome reconstruction, curation, gene-model training, evidence-driven prediction, annotation assembly and quality control without further intervention. As a first step, the pipeline checks the alignment error rate of the reads to avoid using subreads as HiFi reads (Supplementary Note S1). The output is a GFF3 annotation in which every feature carries a tier attribute (*SQANTI_curated* for the curated layer, *Augustus* for the predicted layer), together with the corresponding protein FASTA file and a quality-control report. Following the FAIR principles [28], the workflow is divided into seven modular phases, each of which can be inspected, re-run or replaced independently.

#### Transcriptome reconstruction and curation

Transcript models are reconstructed from the long reads using IsoQuant [10], which was selected for its ability to operate without a reference annotation and for its strong overall performance in LRGASP [13]. When the poly(A) tails of the FLNC reads have been trimmed during pre-processing, IsoQuant is run with the *--polya_trimmed all* option. Alternatively, transcriptome reconstruction can be performed with IsoSeq3 [29], using *isoseq cluster* followed by *collapse*. The resulting transcript models are then curated with SQANTI3 [15].

As SQANTI3 requires a reference annotation, which may not be available for newly assembled genomes, we introduced a *placebo* mode that uses a minimal synthetic annotation. This annotation contains one artificial two-exon transcript in the first kilobase of every sequence of the assembly (chromosome, scaffold or contig), because SQANTI3 processes only the sequences that carry at least one annotated gene; it provides the input required by SQANTI3 without relying on prior transcript structures. Consequently, all transcripts are classified as intergenic and quality control relies on features derived from the genome sequence and the reconstructed transcripts. Filtering is performed using the SQANTI3 rules module, with criteria based exclusively on intrinsic indicators of transcript quality: canonical splice junctions, a predicted coding ORF, a coding sequence (CDS) of at least 100 nt, absence of an intra-priming signature (defined as no more than 59% adenine downstream of the 3′ end) [14,30], absence of predicted nonsense-mediated decay (NMD) and a *PSAURON* coding score of at least 0.7 [31]. Two filtering strategies are available. Under the strict filter, a transcript is retained only if it satisfies all criteria. The flexible filter, applied by default, also retains transcripts that fail a single intra-priming, PSAURON or canonical-junction criterion, provided they are supported by at least ten full-length reads. Both rule sets can be customized to suit the target organism. Because SQANTI3 assigns a distinct novel-gene identifier to each isoform that cannot be mapped to a reference gene, all isoforms are classified as novel relative to the synthetic annotation. Gene identifiers assigned by IsoQuant are therefore restored after filtering to prevent each curated transcript from being incorrectly represented as separate genes in the final annotation. However, IsoQuant can assign isoforms that share splice junctions to different genes, so the numbers of curated genes are upper bounds and the numbers of transcripts per gene lower bounds; transcript-level and locus-level accuracy are not affected.

#### Hybrid gene-model training

Following our previous recommendations [21] and the AUGUSTUS training protocol [32], the workflow uses a default training size of 5,000 genes and a flanking region of 1,000 bp. Both parameters can be adjusted to suit the target organism. The training set is assembled primarily from conserved single-copy orthologs of the user-selected BUSCO lineage [33]. These orthologs are located in the assembly by protein-to-genome alignment with miniprot [34] and retained only if they meet stringent quality criteria, including complete start and stop codons and at least 95% sequence identity. If the selected orthologs do not provide the requested number of genes, the training set is supplemented with the dominant open reading frame from each gene in the curated transcriptome. This hybrid strategy is the default (*mixed*), although users can alternatively train AUGUSTUS using BUSCO orthologs alone (*busco_only*) or curated transcripts alone (*sqanti_only*). BUSCO orthologs provide a consistent training backbone independent of the genes expressed in the lrRNA-seq sample(s) and represent a diverse set of genomic loci that may not be captured by an experiment restricted to a single tissue or condition. Curated transcripts supplement this backbone only when the available orthologs are insufficient to meet the requested training size. AUGUSTUS is then trained with etraining; genes that produce errors in this first round are removed from the training set and the model is retrained.

#### Paralog recovery

lrRNA-seq data provide direct evidence only for the genes expressed in the sequenced sample, leaving unexpressed genes unsupported. To recover additional paralog loci without relying on external protein evidence, which may be unavailable, protein sequences predicted from the curated transcripts are aligned back to the genome using miniprot with a relaxed alignment threshold (*--outs 0.5*). This setting allows secondary alignments to be retained, providing evidence for paralogous loci. By default, miniprot reports only the primary alignment, and does not consider paralog evidence.

#### Hint generation

Intron, CDS, and CDS start and stop hints are derived from the curated transcript models and assigned to the *lrRNA* evidence source. These feature types were selected empirically based on their effectiveness in transferring long-read information into the gene predictor (see Results). Miniprot alignments generated in the paralog-recovery phase are incorporated as a second evidence class, designated as *P*. The supported hint feature types and their corresponding weights, bonuses and penalties for each evidence source are specified in two user-accessible configuration files.

#### Evidence-driven prediction

AUGUSTUS [6,11] uses the species-specific model trained on the combined BUSCO and curated-transcript gene set to perform evidence-driven gene prediction with both classes of hints. AUGUSTUS is run with soft-masking enabled (*--softmasking=1*) and with alternative transcripts from evidence allowed, whereas UTR prediction is disabled. Prediction is performed independently for each contig in *split* mode, enabling parallel execution across the computing cluster.

#### Annotation assembly

The final annotation is assembled by integrating the SQANTI3-curated and AUGUSTUS-predicted gene sets in two tiers. Tier 1 comprises the transcript models curated by SQANTI3, which are incorporated directly into the GFF3 annotation with the attribute *tier=SQANTI_curated*. These models retain their full structure, including CDS and the UTRs when present. Tier 2 comprises AUGUSTUS predictions, identified by the attribute *tier=Augustus*. An AUGUSTUS gene model is retained only if no Tier 1 gene overlaps it on the same strand, giving precedence to SQANTI3 models wherever the two sets overlap.

The inclusion of single-exon AUGUSTUS predictions is controlled by the user-configurable *filter_mode* parameter. Under the *strict* mode, a single-exon prediction is retained only if supported by at least one hint. Under the *medium* mode, the default used in all analyses in this study, it is retained if supported by a hint or if its length is at least 300 bp. Under the *none* mode, all single-exon predictions are retained, irrespective of evidence support or length. Multi-exon predictions are retained without evidence-based filtering in all three modes.

#### Quality control

Annotation quality is assessed using GAQET2 [35,36] and gffcompare [37], both integrated into the SQANTI-evidence pipeline. GAQET2 combines complementary tools to evaluate annotation quality, including AGAT [38] for structural statistics, BUSCO [33] for annotated proteome completeness, and OMArk [39] for proteome consistency. gffcompare is incorporated for benchmarking purposes, as it allows comparison to a reference annotation. To support quality assessment when no reference annotation is available, the pipeline also reports the proportion of reconstructed transcript models classified as non-coding before filtering. This value is contextualized against the range observed across the eight benchmark species, providing a reference for assessing the potential noise level of the input transcriptome.

### Software requirements

SQANTI-evidence v1.1.0 runs on Linux, on a workstation or a Slurm cluster, and requires conda and Snakemake. The benchmark used IsoQuant 3.13, SQANTI3 6.0.2, AUGUSTUS 3.5.0, miniprot, BUSCO 5.8 for the training set, GAQET2 1.15.0 with BUSCO 6.0.0, and gffcompare. BRAKER3 was run from its container and BRAKER4 from commit *42be27b* with the OrthoDB v12 clade partitions.

### Benchmark design

#### Benchmark species, data and accuracy metrics

To evaluate SQANTI-evidence across diverse genomic contexts, we selected eight model organisms with a high-quality reference annotation and publicly available lrRNA-seq data: *Saccharomyces cerevisiae*, *Caenorhabditis elegans*, *Arabidopsis thaliana*, *Drosophila melanogaster*, *Gallus gallus*, *Danio rerio*, *Mus musculus* and *Homo sapiens*. Their genome sizes range from 12 Mb to 3.3 Gb, encompassing a broad range of annotation complexities. The reference genomes were obtained from Ensembl as soft-masked assemblies [40]; for human, the primary assembly was used, without alternate haplotypes. PacBio transcriptomics data were available for seven species. For *D. melanogaster* Nanopore data were used because publicly available Iso-Seq datasets were not available. Dataset accessions are provided in the Availability of data and materials section.

All annotations were evaluated against the coding sequences of the corresponding Ensembl reference annotations. Accuracy was assessed using gffcompare at the transcript and locus levels. A predicted transcript was considered correct at the transcript level if its CDS intron chain matched that of a reference transcript exactly (gffcompare class =). At locus level, a reference gene was considered recovered if at least one predicted transcript matched a reference transcript from that gene exactly. Sensitivity was defined as the percentage of reference transcripts or genes recovered, whereas precision was defined as the percentage of predicted transcripts or loci matching the reference. The F1 score was calculated as the harmonic mean of precision and sensitivity.

We also calculated exact-CDS precision, defined as the proportion of all predicted models whose CDS intron chain matches that of a reference transcript exactly. Because this metric uses all predicted models as its denominator, it differs from the transcript-level precision reported by gffcompare: the two measures are therefore reported separately. To distinguish the contribution of evidence-based curation from that of AUGUSTUS prediction, each run was evaluated twice: first using only the curated layer, and then using the complete integrated annotation.

#### Evaluation of design decisions

To select the transcript reconstruction method, we compared IsoQuant with IsoSeq3 using the same reads from seven species; IsoSeq3 was not evaluated on the Nanopore dataset of *D. melanogaster*. Both methods were assessed on the unfiltered curated layer, allowing reconstruction performance to be compared at transcript and at locus level before any curation.

To evaluate transcript curation, we applied three filter strategies to each species and reconstruction methods: no filtering, the strict filter and the flexible filter. Before running the complete pipeline, candidate rule sets were evaluated directly using the SQANTI3 classification tables obtained from IsoQuant reconstructions. For each rule set, we calculated the proportion of reference-exact isoforms retained and the proportion of retained isoforms that were reference-exact. To identify the conditions responsible for excluding reference-exact isoforms under the strict filter, each excluded isoform was classified according to the condition, or combination of conditions, that it did not meet. Isoforms flagged for intra-priming were grouped by the number of supporting full-length reads, and the proportion of reference-exact isoforms was calculated for each group. The best-performing candidate rule sets were then evaluated through the complete pipeline across all eight species. Finally, we assessed whether the workflow’s reference-free noise diagnostic reflected quality by calculating its Spearman correlation with the proportion of spurious models, as determined using the reference annotations, across the eight species.

To determine the optimal hint types and weights for AUGUSTUS, we ran the final pipeline on human chromosome 19 and on the entire *A. thaliana* genome. We evaluated eight combinations of hint feature types and a control configuration using protein hints alone, together with five weight files. Across these files, the bonus assigned to long-read evidence varied between 10⁴ and 10¹⁵, while the penalty (*malus*) for predictions lacking hint support was reduced below its default value of 1. The trained species-specific model, the curated transcripts, the protein hints and the reference were held constant across configurations. Raw AUGUSTUS predictions were evaluated before integration with the curated layer to isolate the effect of the hint configuration on gene prediction. To quantify the contribution of the predicted layer, the same runs were evaluated using both the complete annotation and the curated layer alone.

To determine how the curated and predicted layers should be combined, we first compared each curated transcript with the corresponding raw AUGUSTUS predictions at the same genomic position, before layer integration, across all eight species. For each curated transcript, we assessed whether AUGUSTUS reproduced its CDS intron chain, and evaluated both models against the reference annotation. We then evaluated all AUGUSTUS models discarded by the integration rule, classifying them according to whether their intron chain was already present in the curated layer, matched a reference transcript despite being absent from the curated layer, or matched no reference transcript at all. Single-exon predictions were analyzed separately. Finally, we tested an additional evidence requirement for the predicted layer, under which multi-exon models lacking a protein-supported intron were removed. We measured the effect of this filter on exact-CDS precision and on the number of reference-exact models retained.

#### Comparison with BRAKER3 and BRAKER4

Our eight benchmarking species were also annotated with BRAKER3 [19] and BRAKER4 [20]. Both methods follow an evidence-driven approach similar to that of SQANTI-evidence, and differ in the way they use protein evidence: BRAKER3 accepts user-supplied proteins whereas BRAKER4 selects proteins from OrthoDB [41] according to the target clade. BRAKER3 was run in long-read mode using the same reads as in SQANTI-evidence, and unfiltered protein sequences inferred from the reconstructed transcripts. BRAKER4 was run with default settings and the OrthoDB v12 clade partition, representing a standard protein-supported annotation workflow. GeneMark-ETP [42] failed reproducibly on *S. cerevisiae* with both BRAKER versions, so comparisons were restricted to the remaining seven species. For this comparison, SQANTI-evidence was run with IsoQuant and the flexible filter. We evaluated accuracy with gffcompare and assessed completeness of the annotations with GAQET2, using BUSCO genes, OMArk consistency and missing genes. We also compared the total numbers of genes, transcripts and transcripts per gene.

To investigate differences in locus sensitivity, reference protein-coding genes were divided into two groups according to whether they overlapped at least one IsoQuant model before filtering. Genes overlapping at least one model were classified as having read evidence, whereas those with no overlap were classified as lacking evidence. Locus sensitivity was calculated separately for each group.

To investigate the differences in precision, the curated and predicted layers of SQANTI-evidence were evaluated separately against both BRAKER annotations using exact-CDS precision. For each predicted model, we counted the number of introns supported by protein hints, and classified the models as having all introns supported, being a multi-exon model without protein support, or being a single-exon model. The same classification was applied to the AUGUSTUS-derived models retained in the final BRAKER annotations by TSEBRA [22], the transcript selector of BRAKER, which removes models with insufficient evidence support; the AUGUSTUS predictions discarded by TSEBRA were not evaluated. Non-exact predicted models were further classified by genomic context: overlap with a non-coding gene or pseudogene, overlap with a coding gene of a different structure, location in unmasked sequence without an annotated gene, or predominant location within soft-masked sequence.

Because each SQANTI-evidence model carries a tier attribute, alternative annotation subsets can be generated after the workflow has completed. To assess the effect of such filtering on annotation precision, we defined five subsets: the default annotation (A); the predicted layer restricted to models whose introns are all supported by protein hints (B); subset B excluding single-exon predicted models (C); the curated layer alone (D); and subset D with curated loci that do not overlap any annotated gene excluded from the precision denominator (E). Subsets A and D are selected with the tier attribute, B and C additionally require the protein-hint file written by every run (Supplementary Note S3), and E requires a reference annotation.

To assess isoform recovery, we selected reference genes with at least two annotated CDS isoforms and counted the number for which each annotation recovered at least one or at least two isoforms exactly.

Untranslated region (UTR) annotation was evaluated on transcript models whose CDS were predicted exactly by both SQANTI-evidence and BRAKER4. For each end at which the reference annotation included a UTR, we recorded whether the method annotated a UTR at that end (coverage) and, when present, the distance between the annotated and the reference transcript end (error). Distances were signed, with negative values indicating that the predicted end fell within the reference transcript and positive values indicating that it extended beyond the reference end. In SQANTI-evidence, UTR boundaries correspond to the ends of the full-length reads in the curated layer. In BRAKER4, UTRs are produced by stringtie2utr from a StringTie assembly [9] generated using the same reads. BRAKER3 does not predict UTRs and was therefore excluded from this analysis.

#### Annotation of ERGA species

To evaluate the applicability of SQANTI-evidence to newly assembled genomes, we obtained six assemblies and their corresponding long-read datasets from the ERGA pilot [5]: *Xylocopa violacea* (violet carpenter bee), *Alopecurus aequalis* (foxtail grass), *Bufotes viridis* (green toad), *Perca* sp. ‘yellow fin Alpine’ and *Zingel asper* (percid fishes), and *Salvelinus alpinus* (Arctic char). Genome and read accessions are provided in the Availability of data and materials section; the accessions of *Perca* and *Zingel* were still under review at ENA at the time of writing. None of the six assemblies had been repeat-masked on delivery. Each genome was therefore annotated in its original form and after soft-masking with Red [43], allowing us to assess the effect of repeat masking on annotation outcomes.

We additionally annotated seven ERGA invertebrate datasets sequenced by Martínez-Redondo *et al.* [25]. These datasets comprised two gastropods (*Phorcus turbinatus*, *Physella acuta*), a polychaete (*Arenicola marina*), a planarian (*Schmidtea mediterranea*), a nemertean (*Tetrastemma melanocephalum*, two libraries) and one nematode (*C. elegans*). The corresponding genome assemblies were provided with 31–56% of their sequence soft-masked. The *C. elegans* dataset was sequenced at 34-fold lower depth than the benchmark dataset for this species and was used as a sequencing-depth control rather than as a new annotation. All ERGA genomes were annotated with IsoQuant for transcriptome reconstruction and the SQANTI3 flexible filter. Because GAQET2 does not provide a suitable BUSCO lineage for *B. viridis*, annotation completeness was assessed using *tetrapoda_odb12*, the closest available lineage.

## Declarations

### Ethics approval and consent to participate

Not applicable.

### Consent for publication

Not applicable.

### Availability of data and materials

SQANTI-evidence is available on GitHub [44] under the GPL-3.0 license. The version used in this study (v1.1.0, commit *23b7d75*, released 17 September 2026) is archived on Zenodo [45] (concept DOI 10.5281/zenodo.22682045; version DOI 10.5281/zenodo.22816381). Every run reported here is reproducible with this version, and a confirmation run on yeast with the tagged code reproduced the gene models of the benchmark run exactly. The annotations of the 13 ERGA datasets, with their protein and CDS sequences and their GAQET2, BUSCO and OMArk reports, are available on Zenodo (https://doi.org/10.5281/zenodo.23057110) [46]. The analysis code, run configurations, derived tables and figure scripts, together with the final annotations, protein sets and gffcompare and GAQET2 outputs of every benchmark run and species, are available in the benchmark repository on Zenodo (https://doi.org/10.5281/zenodo.23086233) [47].

The datasets analyzed in this study are available from public repositories under the following accessions (Table 1). *H. sapiens*: WTC11, LRGASP FLNC [13], nine ENCODE runs merged (ENCFF112MRR, ENCFF122GKS, ENCFF338WQL, ENCFF434SWA, ENCFF455RXJ, ENCFF620NFT, ENCFF875XMU, ENCFF962OWJ, ENCFF992WSK); *M. musculus*: brain and kidney Iso-Seq, ERR14205062 (PRJEB85167) and ERR15371799 (PRJEB94912) [48]; *G. gallus*: ERR7015103 (PRJEB48060, GENE-SWitCH), HiFi regenerated with *ccs*; *D. rerio*: SRR32588721 (PRJNA1232602); *D. melanogaster*: SRR29784392, SRR29784417, SRR29784414 (PRJNA1134728, Nanopore); *A. thaliana*: ERR15191386 (PRJEB91362); *C. elegans*: ERR17585088–90 (PRJEB121150, N2 Revio); *S. cerevisiae*: SRR21679915, SRR23265078, SRR21679893, SRR23265077 (PRJNA883530). Genomes and annotations are from Ensembl release 115 [40] (TAIR10 and Ensembl Plants 62 for *Arabidopsis*). ERGA pilot data: *Xylocopa violacea* GCA_963969225.2, PRJEB72102 (ERR13064450–53); *Bufotes viridis* GCA_963931945.2, PRJEB71764 (ERR12542608); *Alopecurus aequalis* GCA_964340505.1, PRJEB75647 (ERR13928836, ERR13928839); *Salvelinus alpinus* GCA_964197945.1, PRJEB76174 (ERR13703899, ERR13703946–48); *Perca* sp. ‘yellow fin Alpine’ fPerYfa1 (PRJEB122538), PRJEB107436 (SAMEA122781568; ERR18007935–47); *Zingel asper* fZinAsp1.1 (PRJEB122083), PRJEB107293 (SAMEA122719616; ERR16908784, ERR16908787, ERR16908789, ERR16908791, ERR16908793, ERR16908795); the *Perca* and *Zingel* accessions were under review at ENA at the time of writing. ERGA invertebrate Iso-Seq data from Martínez-Redondo et al. [25], ENA umbrella project PRJEB114751: *Phorcus turbinatus* GCA_965237135.1, PRJEB108696 (ERR17004618–19); *Physella acuta* GCF_028476545.2, PRJEB108697 (ERR17004620–21); *Arenicola marina* GCA_964019135.1, PRJEB108691 (ERR17004611); *Schmidtea mediterranea* GCA_057920485.1, PRJEB108587 (ERR17004635); *Tetrastemma melanocephalum* GCA_977011905.1, PRJEB108704 (ERR17004629, ERR17004630); *C. elegans* GCF_000002985.6 (WBcel235), PRJEB108700 (ERR17004624).

## Competing interests

AC has received in-kind funding from Pacific Biosciences for library preparation and sequencing. AC has collaborated with Oxford Nanopore in the Marie Skłodowska-Curie Actions Doctoral Network project LongTREC. PAL declares no competing interests.

## Funding

This work was supported by the OSCARS project, which has received funding from the European Commission’s Horizon Europe Research and Innovation programme under grant agreement No. 101129751; by the Spanish Ministerio de Ciencia, Innovación y Universidades (MICIU) under project **[MICIU project code]**; and by the CSIC Conexión Biología Computacional y Bioinformática (Conexión BCB) **[code]**, which funded a research visit to Seville. The funders had no role in the design of the study, the analyses or the writing of the manuscript.

## Authors’ contributions

PAL designed and implemented the workflow, performed the benchmark and the ERGA annotations, and wrote the manuscript. AC conceived and supervised the study and revised the manuscript. All authors read and approved the final manuscript.

## Acknowledgements

The computations were performed on the high-performance computing cluster Garnatxa of the Institute for Integrative Systems Biology (I2SysBio), a joint center of the University of Valencia and the Spanish National Research Council (CSIC). We thank Aureliano Bombarely and Víctor García-Carpintero for their help in understanding GAQET2 and integrating it into SQANTI-evidence, and the ERGA data providers, in particular Rosa Fernández, Carlos Vargas and Laura Triginer, for access to the long-read datasets.

## Notes

https://github.com/ConesaLab/SQANTI_evidence

https://doi.org/10.5281/zenodo.22682045

https://doi.org/10.5281/zenodo.23057110

https://doi.org/10.5281/zenodo.23086233

### Competing Interest Statement

The authors have declared no competing interest.

