## supplementary materials for "SQANTI-evidence: isoform-resolved, evidence-tiered structural annotation of unannotated genomes from long-read RNA-seq"

### Supplementary material

#### *Supplementary figures*

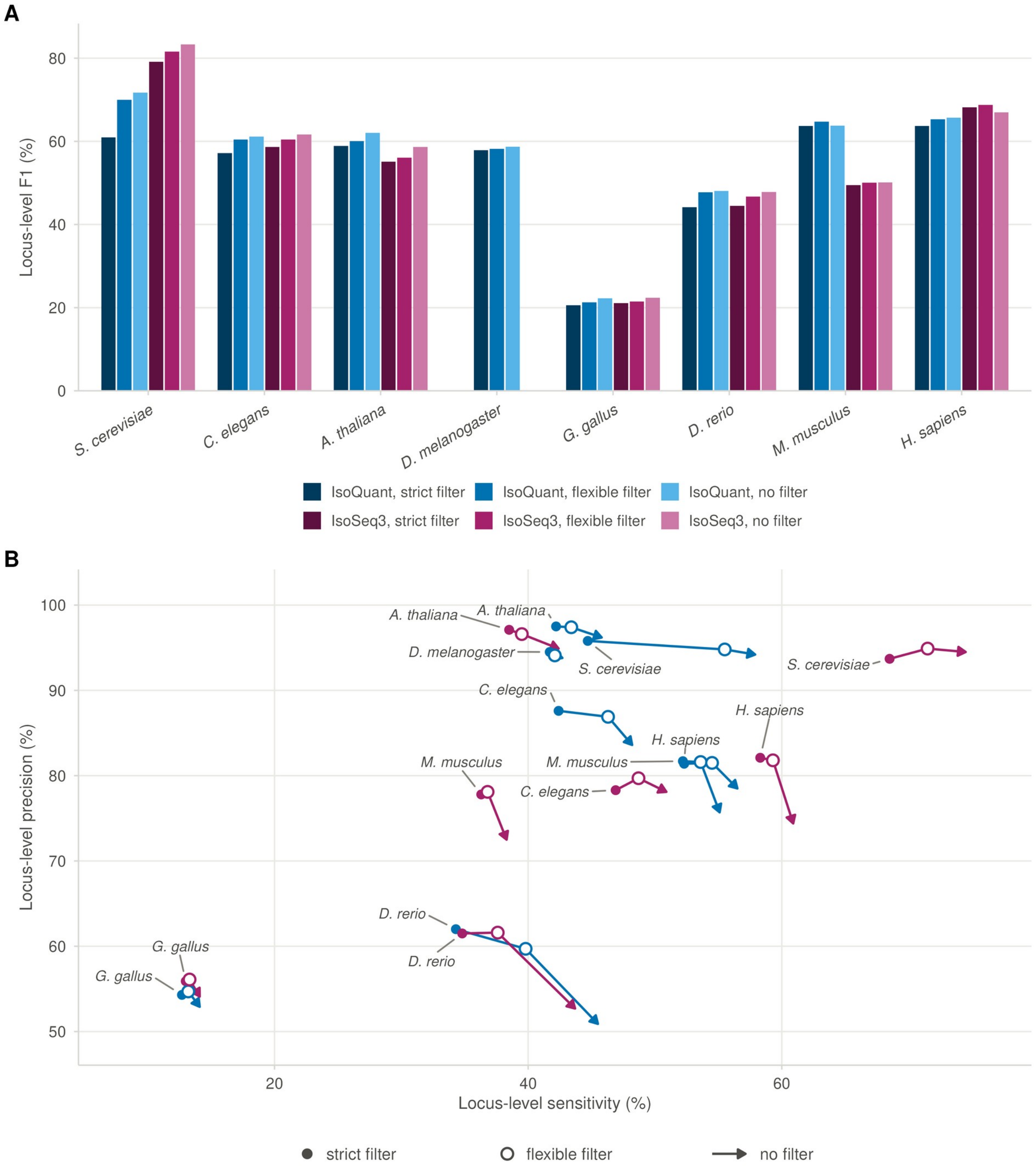


**Figure S1 | Reconstruction method and curation filter at locus level.** The runs of Figure 2 scored on Tier 1 at CDS locus level. A, Locus-level F1. B, Locus-level sensitivity against precision, with the same symbols as Figure 2B. Without a filter, the two reconstruction methods are within four points of locus-level F1 of each other in five species.

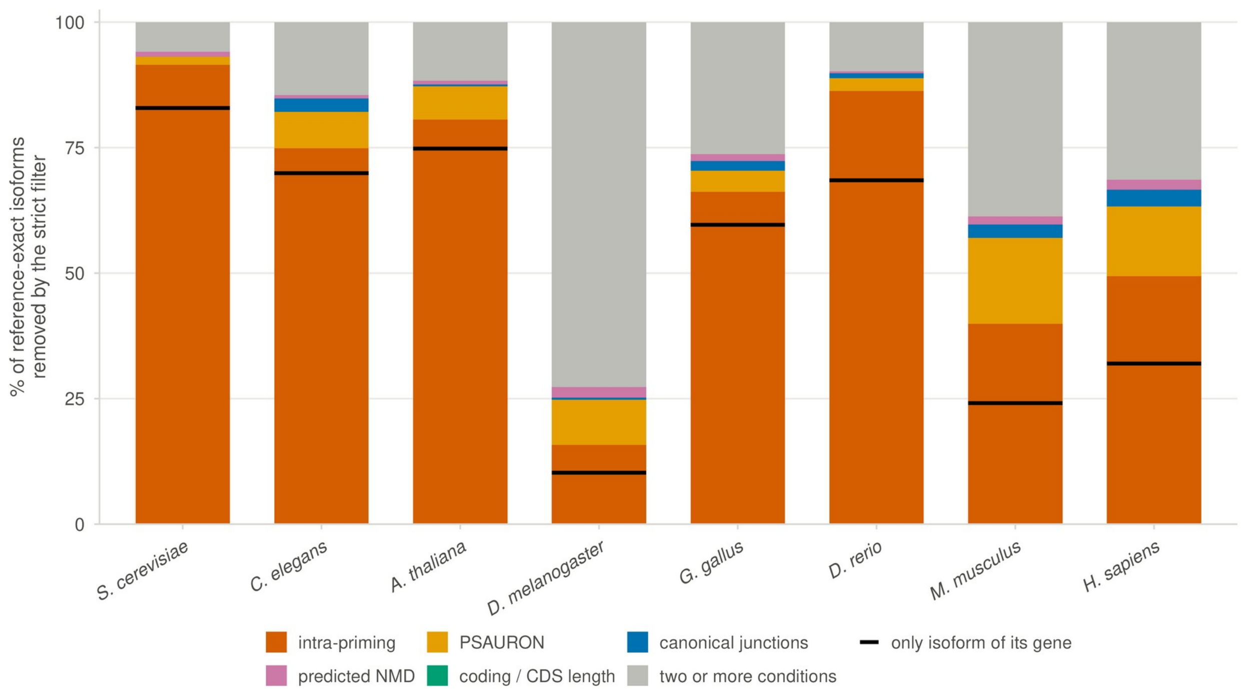


**Figure S2 | The intra-priming condition causes most of the loss of reference-exact isoforms under the strict filter.** For each species, the reference-exact isoforms of the IsoQuant reconstruction that the strict filter removes, split by the condition they fail (intra-priming, PSAURON coding score, canonical junctions, predicted NMD, coding or CDS length), each counted only where it is the sole failing condition; the remainder failed two or more conditions at once. The black line on each intra-priming bar marks the isoforms that were the only isoform of their gene, so that the whole gene left the curated layer; these were 60–93% of the isoforms lost to intra-priming alone, lowest in mouse and highest in worm. Eight species. The intra-priming condition alone accounts for 66–92% of the loss in five of the eight species.

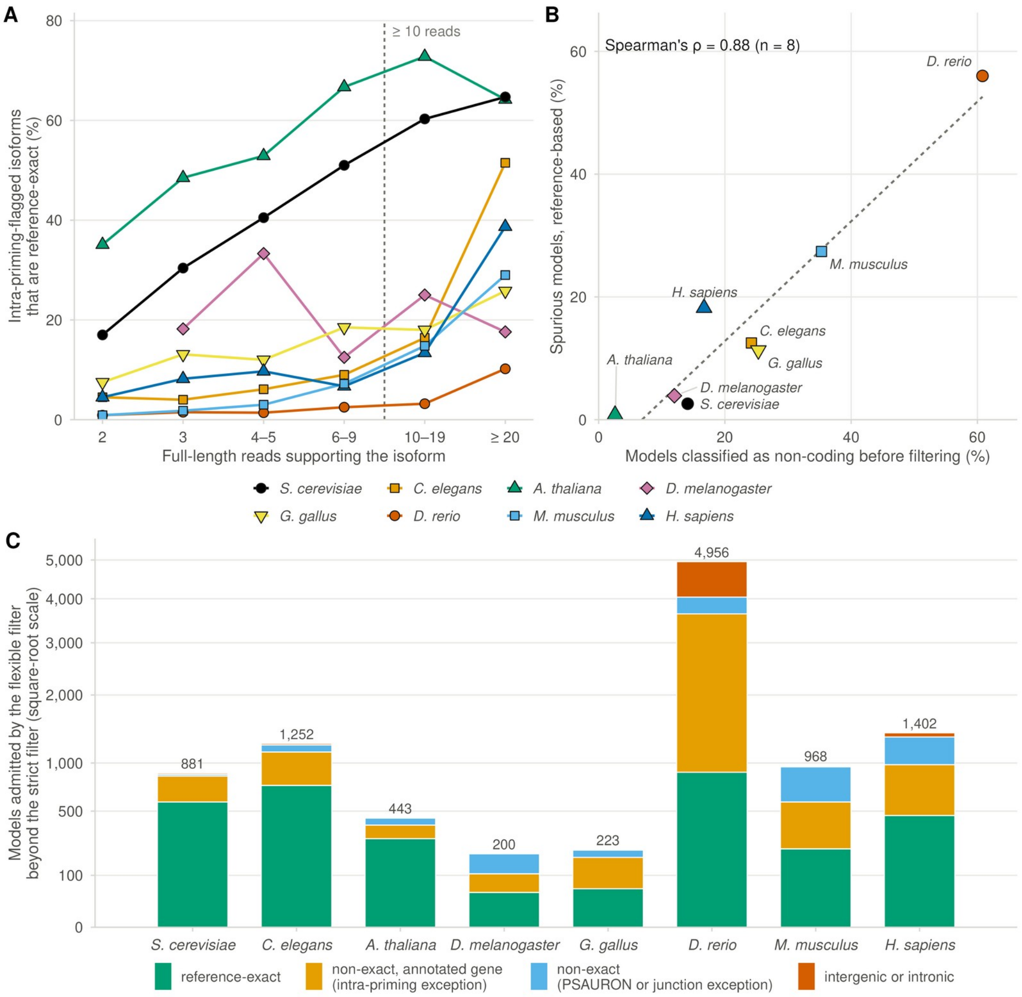


**Figure S3 | Read support, a reference-free noise diagnostic and the composition of the admitted models.** Isoform-level analysis of the SQANTI3 classification tables of the IsoQuant reconstruction before prediction and layer combination, eight species (colors and symbols as in the key). **A**, Isoforms flagged by the intra-priming condition that match a reference transcript exactly, as a percentage of the flagged isoforms in each bin of supporting full-length reads; the dashed line marks the ten-read threshold of the flexible filter. **B**, The noise diagnostic reported by the workflow before filtering: the percentage of reconstructed models classified as non-coding against the reference-based percentage of spurious models, one point per species (n = 8); Spearman's ρ is given in the panel and the dashed line is the least-squares fit. **C**, Models admitted by the flexible filter beyond the strict filter, by class: reference-exact; non-exact isoforms of an annotated gene admitted through the intra-priming exception; non-exact models admitted through the PSAURON or canonical-junction exception; intergenic or intronic. The number above each bar is the total; the y axis is square-root scaled so that the 4,956 admissions in zebrafish do not flatten the 200–1,402 of the other species. In every species except the Nanopore fly dataset the exact fraction of flagged isoforms rises with read support, the non-coding fraction tracks the spurious fraction across species, and only in zebrafish do intergenic or intronic admissions (916) match the reference-exact ones (891).

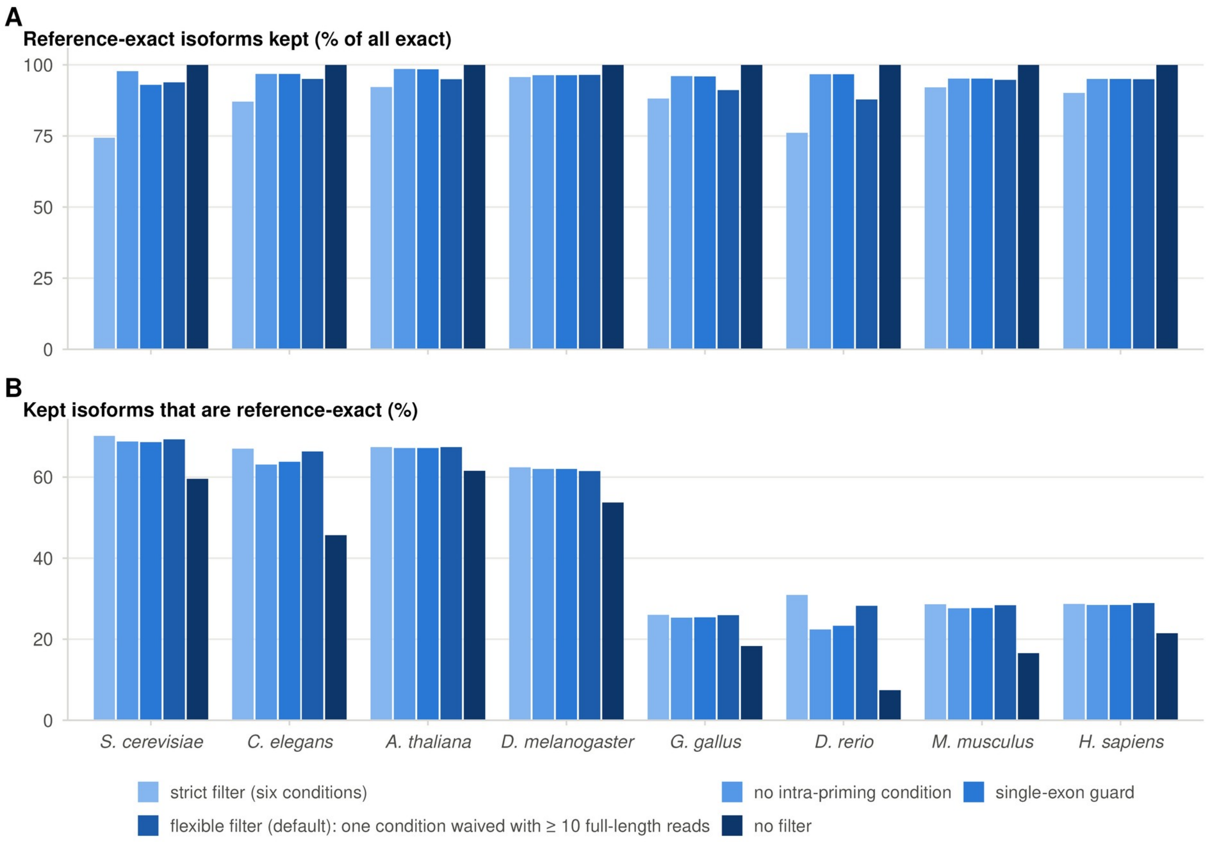


**Figure S4 | The flexible filter recovers most reference-exact isoforms lost by the strict filter.** Candidate rule sets scored on the SQANTI3 classification tables of the IsoQuant reconstruction before prediction and layer combination, eight species: the strict filter (six conditions), three alternatives (no intra-priming condition; a single-exon guard, with no intra-priming condition for spliced models and a relaxed threshold of 79% adenine for single-exon models (Supplementary Note S2); and the flexible filter, the default, in which a model that fails one condition (intra-priming, coding potential or canonical junctions) is kept when it has ten or more full-length reads) and no filter. **A,** Reference-exact isoforms kept, as a percentage of all reference-exact isoforms. **B,** Kept isoforms that are reference-exact. Removing the filter altogether costs precision in every species.

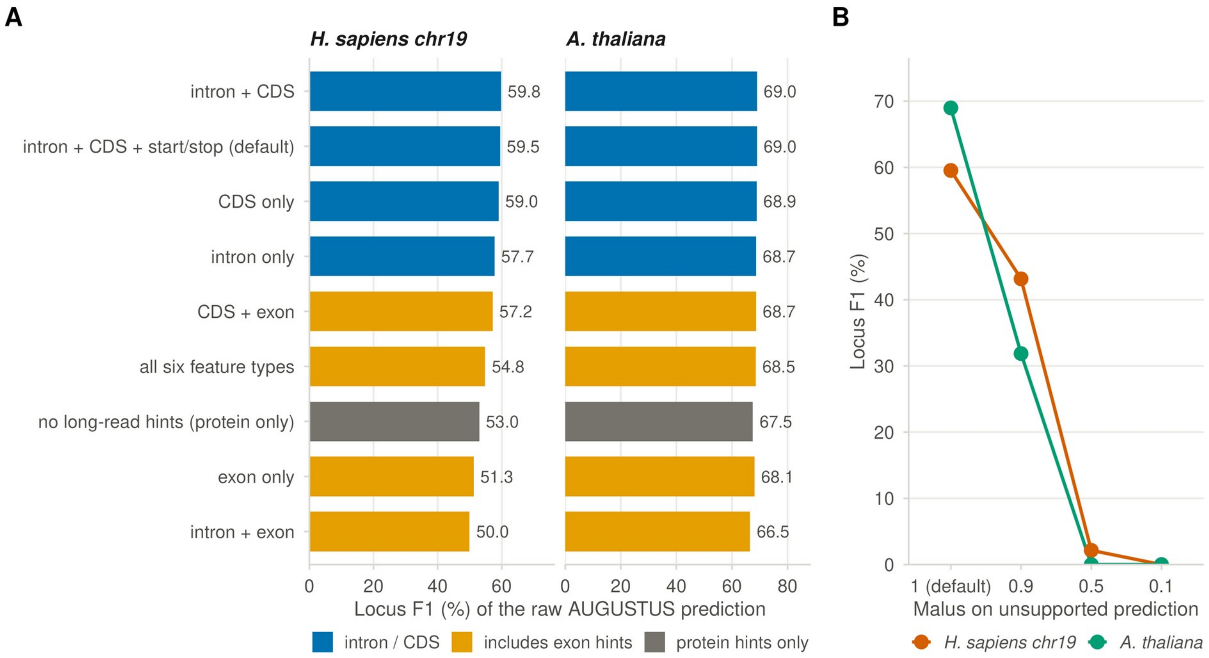


**Figure S5 | Hint feature types matter for AUGUSTUS; a malus below 1 degrades the ab initio prediction.** Raw AUGUSTUS prediction scored against the reference CDS before the two layers were combined, on human chromosome 19 and the whole *A. thaliana* genome, with the trained model, the Tier 1 transcripts, the protein hints and the reference held fixed. A, Locus F1 for nine hint feature sets; the protein-hints-only configuration is the control. Sets that include exon hints and no CDS hints fall below the protein-only control in human. B, Locus F1 as the malus on unsupported prediction is lowered from 1 (default) to 0.1, bonuses at their default values.

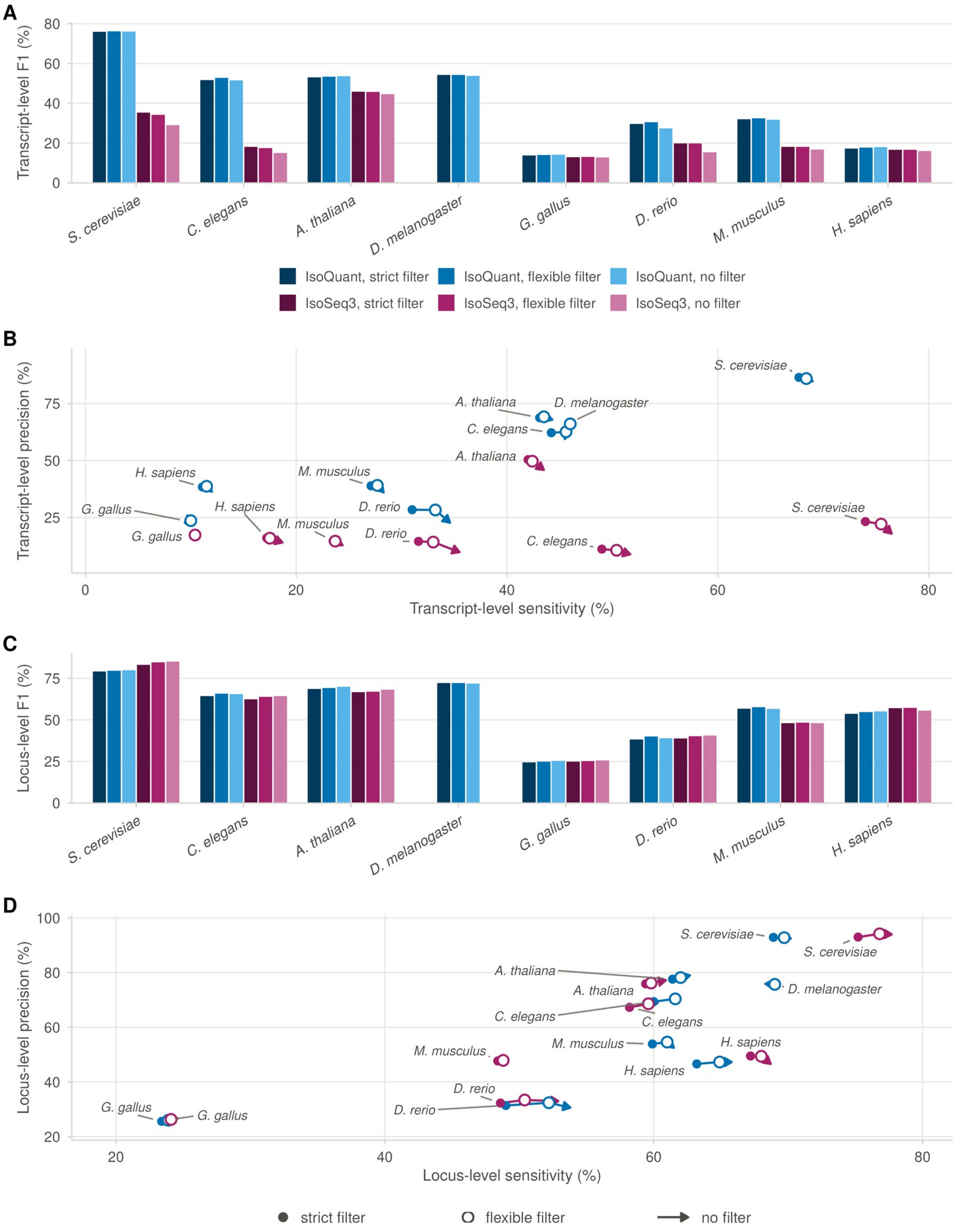


**Figure S6 | Adding the predicted layer compresses the filter effect but not the reconstruction ranking.** The runs of Figure 2 scored on the whole annotation (Tier 1 plus Tier 2). A, B, transcript level; C, D, locus level; symbols as in Figure 2. With IsoQuant, the flexible filter leads the strict filter by −0.1 to 1.1 points of transcript F1 instead of 0.3–8.4 on Tier 1 alone.

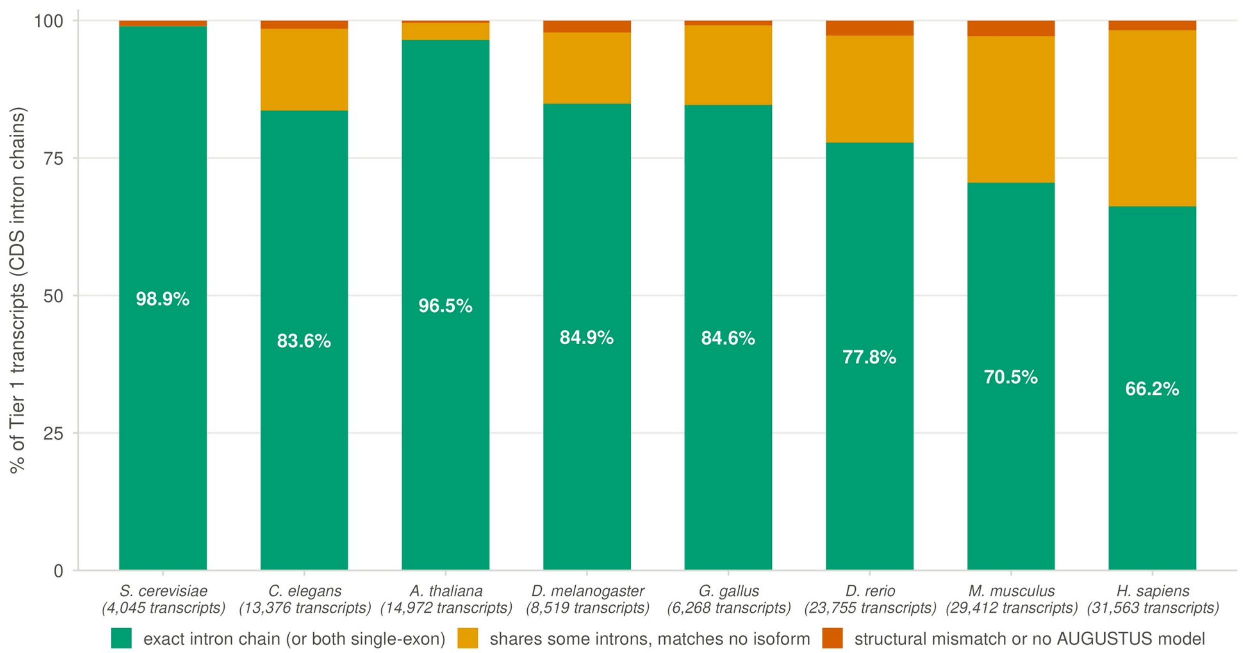


**Figure S7 | AUGUSTUS misses part of the observed CDS structure at 30–34% of mammalian curated transcripts.** Every curated (Tier 1) transcript of the flexible-filter run matched to the raw AUGUSTUS prediction at the same genomic position before the two layers were combined, eight species; the comparison uses CDS intron chains only, so that UTR introns, which AUGUSTUS does not predict without UTR mode, do not count against it. Transcripts per species under the names. Green, AUGUSTUS reproduces the observed CDS intron chain or agrees that the CDS is single-exon (value shown); orange, it shares some CDS introns with the observed transcript but matches no observed isoform; red, structural mismatch or no AUGUSTUS model. Agreement exceeded 96% in yeast and *Arabidopsis*, was 78–85% in worm, fly, chicken and zebrafish, and fell to 70.5% in mouse and 66.2% in human, where the shortfall was mostly a partial match (26.6% and 32.0% of transcripts) rather than a structural mismatch or a missing model (no more than 2.9% in any species). The comparison of both model sets against the reference at the same loci is in Table S3. When UTR introns were included in the comparison, agreement fell to 28.2–70.9% outside yeast; the transcripts that differed from the AUGUSTUS model only by introns in their UTRs accounted for 14.5% of curated transcripts in worm, approximately a quarter in fly and Arabidopsis, 31.9% in zebrafish and 37.3–38.2% in chicken, mouse and human.

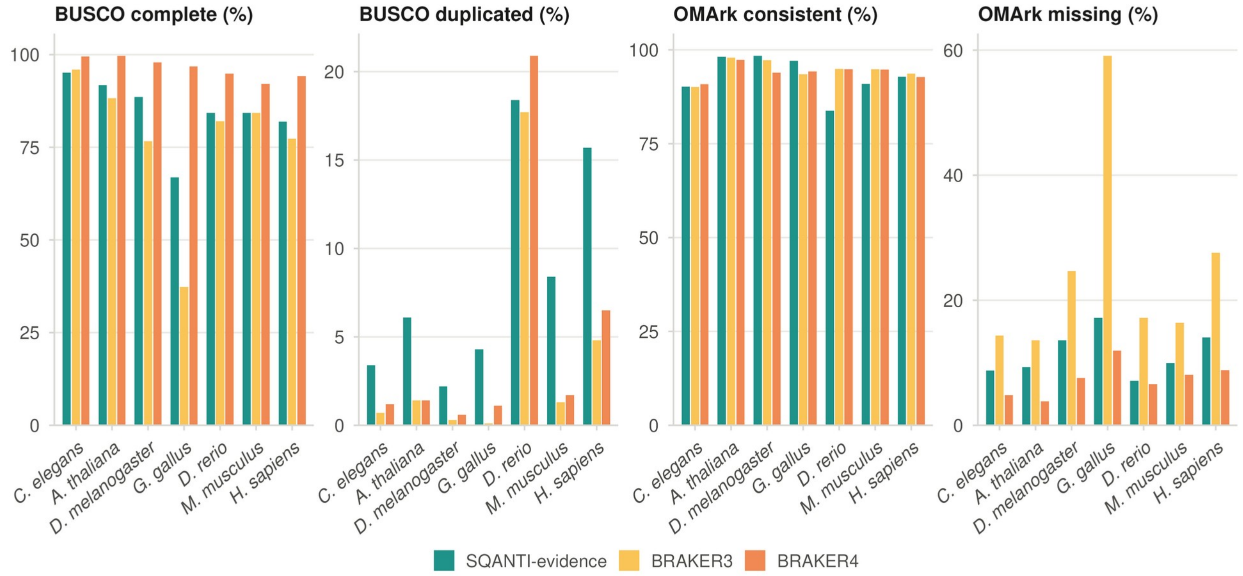


**Figure S8 | GAQET2 completeness, duplication and consistency of the three annotations.** For each species, the percentage of complete and of duplicated BUSCOs and the percentage of OMArk-consistent and of OMArk-missing proteins in the protein sets of SQANTI-evidence (flexible filter), BRAKER3 (proteins from the same long reads) and BRAKER4 (OrthoDB v12); colors as in Fig. 4. Seven species; *S. cerevisiae* is omitted because it has no BRAKER run. BUSCO lineage sets (OrthoDB v12) and OMArk ancestral levels are those chosen by GAQET2 for each species (Methods). Fig. 4A shows BUSCO completeness, OMArk missing genes and gene counts for the same annotations; this figure adds the duplicated BUSCOs and the OMArk-consistent fraction. SQANTI-evidence recovered more complete BUSCOs than BRAKER3 in five of the seven species and had fewer OMArk-missing genes in all seven, but it also contained more duplicated BUSCOs in every species and, in zebrafish, mouse and human, a smaller OMArk-consistent fraction; BRAKER4 was the most complete annotation on both completeness measures in every species.

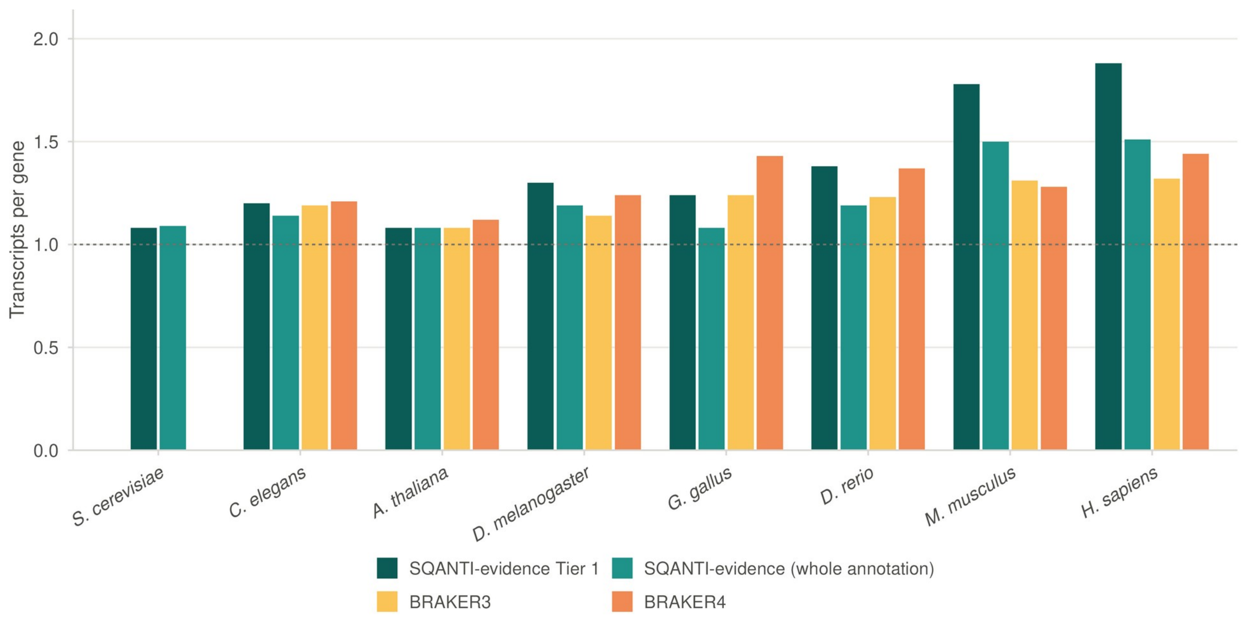


**Figure S9 | Transcripts per gene range from 1.08 to 1.51 in every whole annotation and are highest in Tier 1 of the mammals.** Tier 1 alone, the whole SQANTI-evidence annotation and the two BRAKER annotations, eight species; the dashed line marks one transcript per gene. Tier 2 (not shown) carries 1.01–1.12 transcripts per gene and the BRAKER annotations 1.08–1.44; Tier 1 reaches 1.78 in mouse and 1.88 in human.

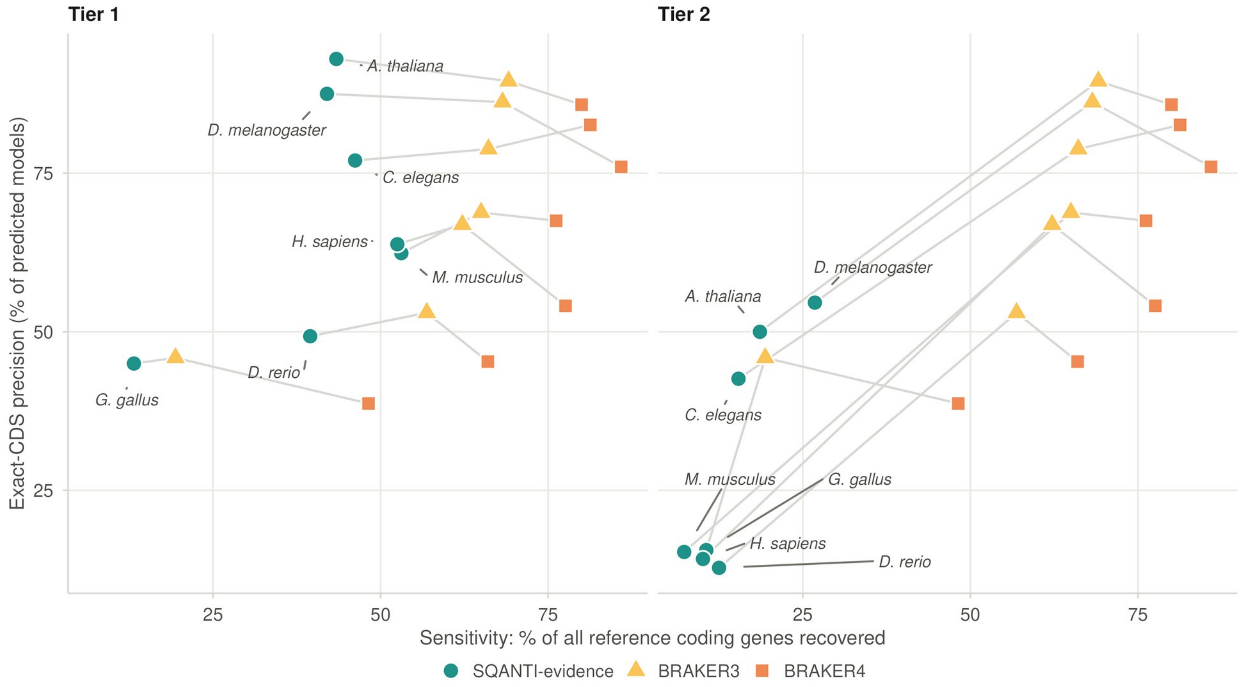


**Figure S10 | Tier 1 provides precision and Tier 2 adds sensitivity at a precision cost, per model.** Sensitivity on all reference coding genes against exact-CDS precision, the percentage of predicted models whose CDS intron chain matches a reference transcript exactly, for Tier 1 (left) and Tier 2 (right) of the flexible-filter run; the BRAKER3 and BRAKER4 points repeat in both panels and gray segments join the three annotations of a species. On this axis Tier 1 exceeds BRAKER4 in five of seven species and trails BRAKER3 in five; Tier 2 is 13–55% exact.

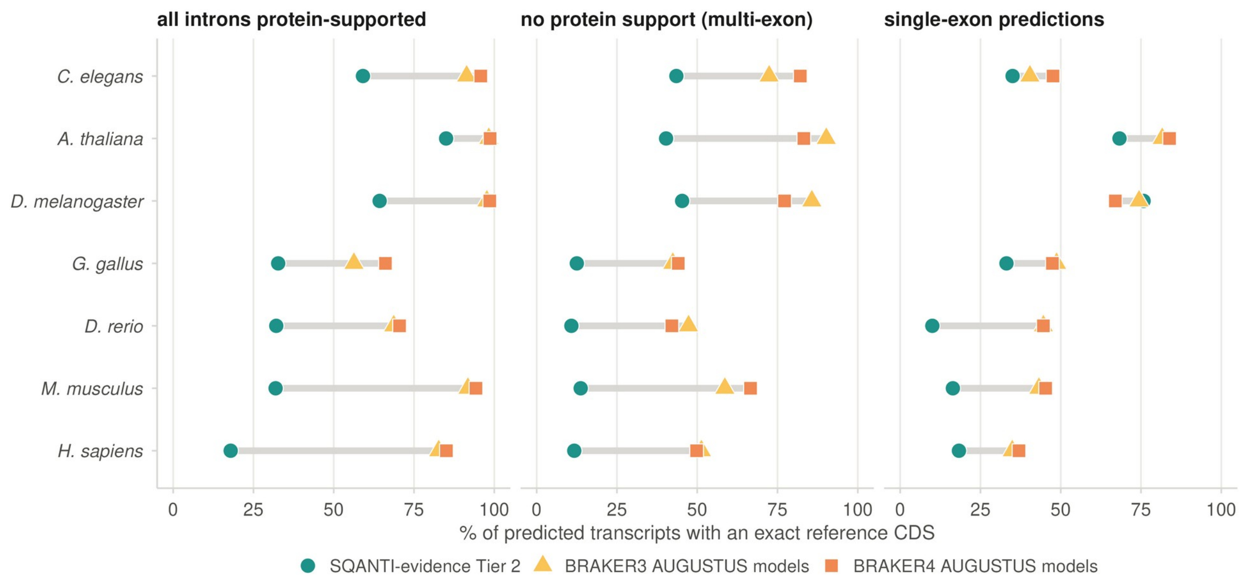


**Figure S11 | Tier 2 is less exact than the AUGUSTUS models BRAKER retains, in all three support classes, except single-exon models in *D. melanogaster*.** Percentage of predicted transcripts with an exact reference CDS for models whose introns are all protein-supported, multi-exon models without protein support, and single-exon models: SQANTI-evidence Tier 2 against the AUGUSTUS-derived models that TSEBRA retained in the final annotation of BRAKER3 and BRAKER4, seven species. The AUGUSTUS output of BRAKER before TSEBRA selection was not scored. Without protein support Tier 2 is 40–45% exact in the compact genomes and 11–14% in the vertebrates.

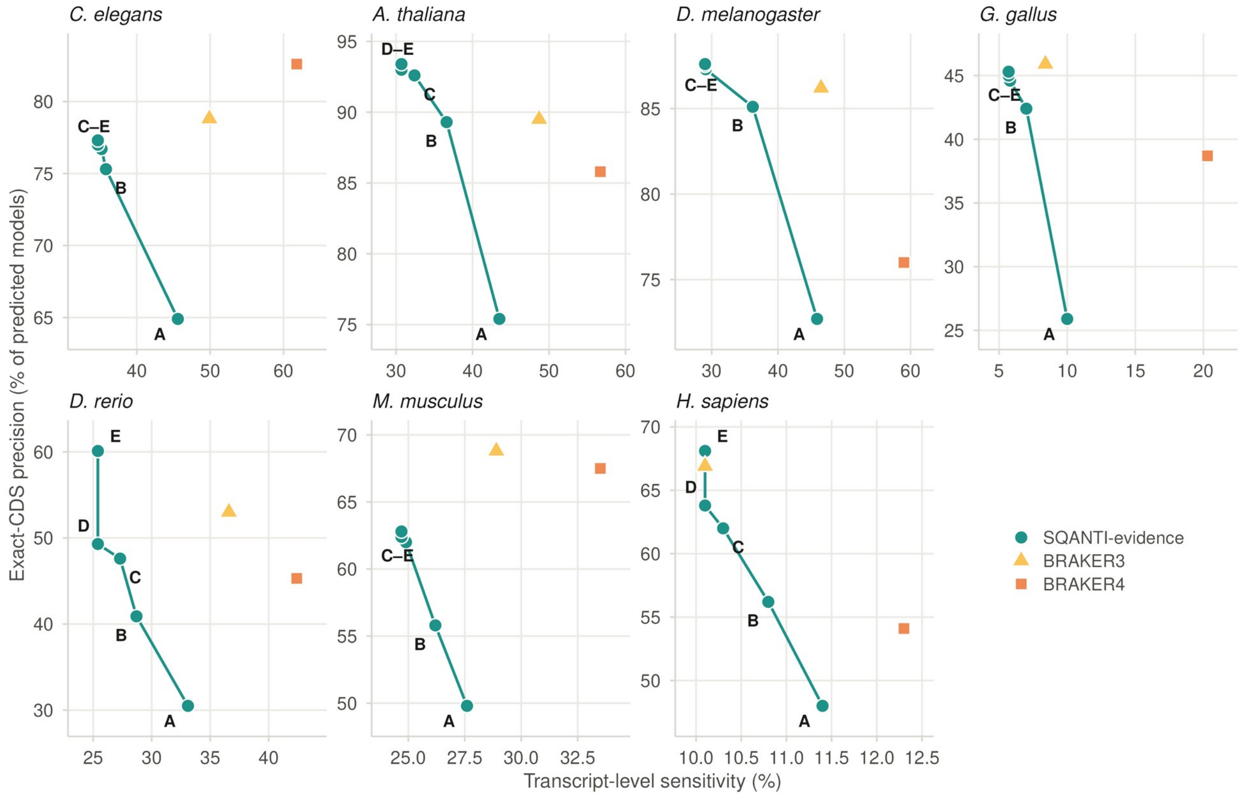


**Figure S12 | Subsets of the SQANTI-evidence annotation at transcript level.** The scenarios of Figure 5 (A, default annotation; B, Tier 2 restricted to fully protein-supported models; C, B without single-exon Tier 2 models; D, Tier 1 alone; E, D with Tier 1 loci overlapping no annotated gene excluded from the precision denominator) for the seven species with BRAKER runs, one panel per species with its own axes, with BRAKER3 (triangles) and BRAKER4 (squares) for the same species; steps that fall on the same point share a label. Sensitivity is the percentage of reference CDS transcripts matched exactly (gffcompare class =) by at least one predicted model, each reference transcript counted once, computed in the same way for SQANTI-evidence and BRAKER. Precision is the percentage of predicted models whose CDS intron chain matches a reference transcript exactly (per model, as in Table 2 and Figure S10). Tier 1 alone (D) is more precise than both BRAKER annotations in *A. thaliana* and *D. melanogaster* and more precise than BRAKER4 in five of the seven species, but less precise than BRAKER3 in *C. elegans*, *G. gallus*, *D. rerio*, *M. musculus* and *H. sapiens*.

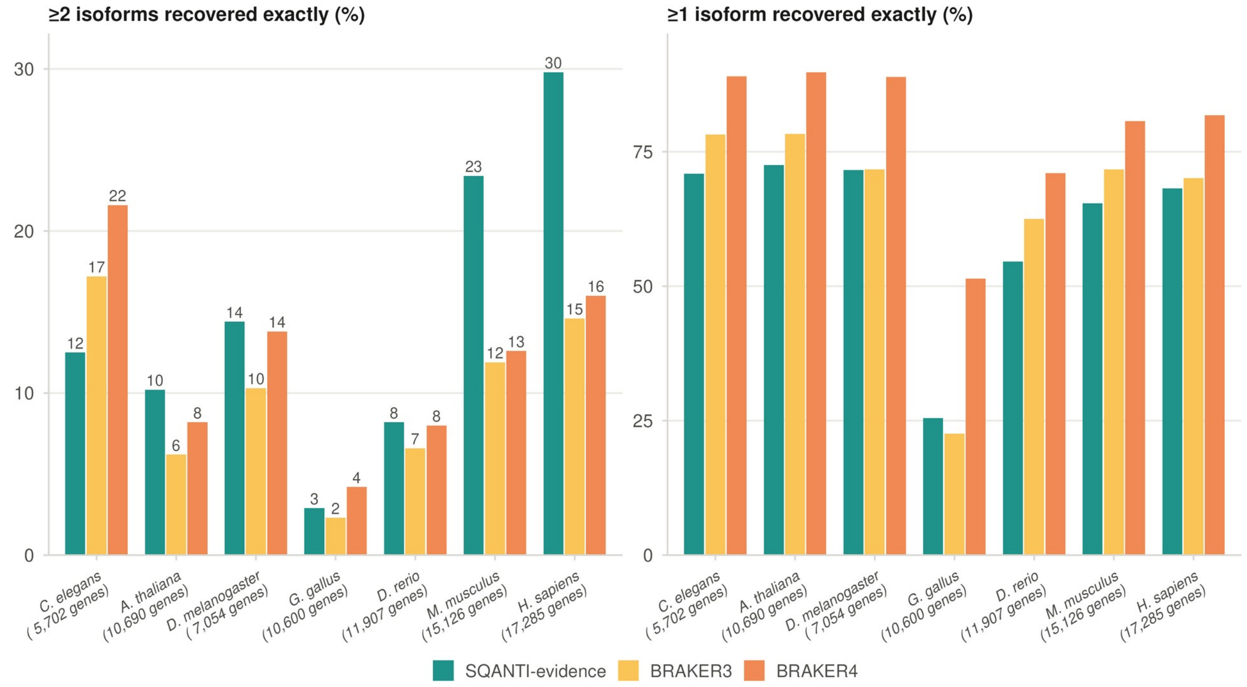


**Figure S13 | SQANTI-evidence recovers multiple exact isoforms per gene twice as often as BRAKER in mammals.** Reference genes with two or more annotated CDS isoforms (count under each species): percentage recovered with at least two exact isoforms (left, values on the bars) and with at least one (right), for the three annotations in seven species. BRAKER4 recovers more genes with at least one exact isoform everywhere; SQANTI-evidence is behind both BRAKER annotations on both counts only in worm, and behind BRAKER4 on both counts also in chicken.

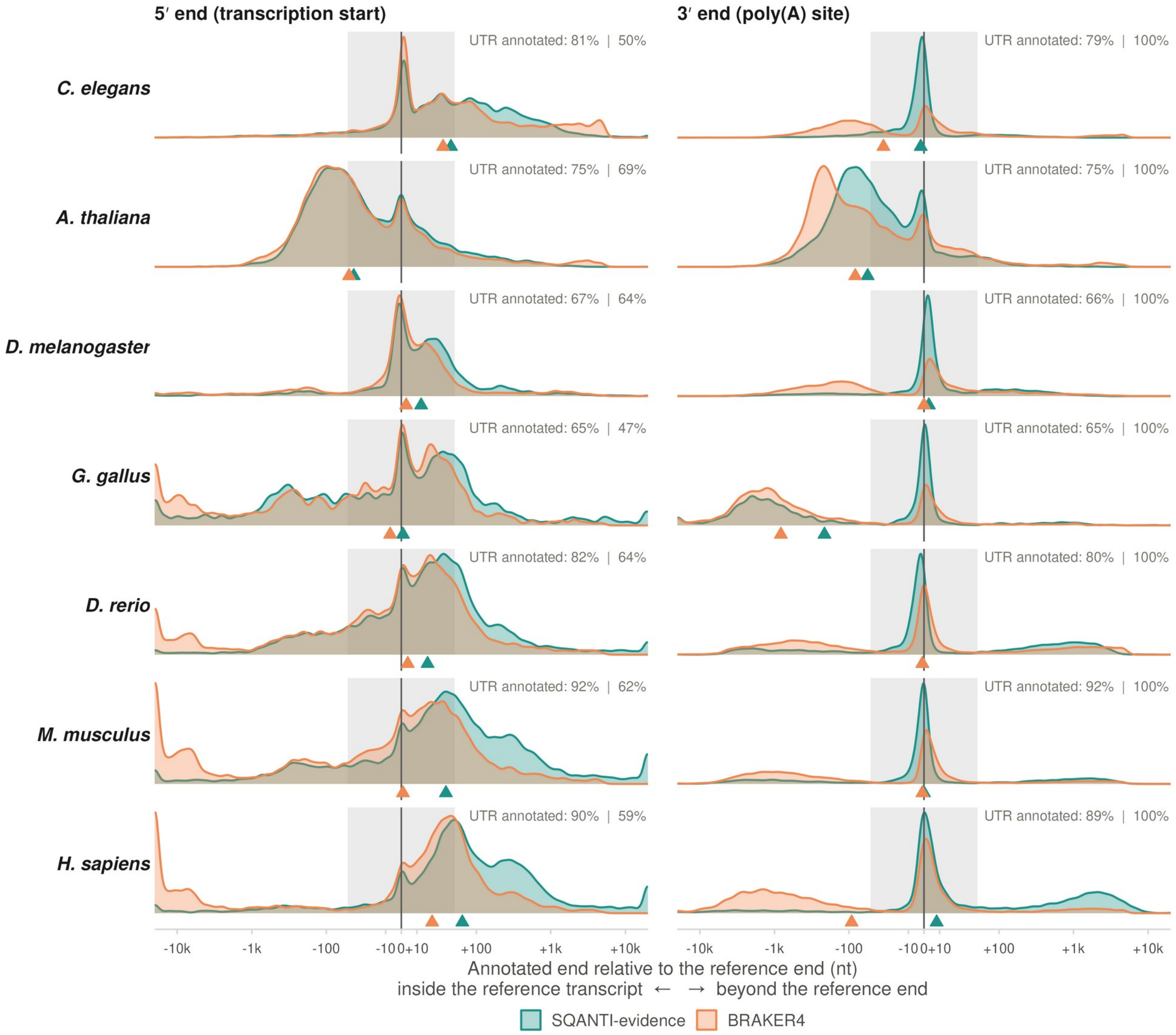


**Figure S14 | Curated 3′ ends lie on the poly(A) site and 5′ ends extend beyond the start.** Distribution of the signed distance from the annotated end to the reference end for every scored end of Figure 6 (negative, the annotated end lies inside the reference transcript; positive, beyond it), SQANTI-evidence and BRAKER4, seven species, on a symmetric logarithmic axis; ends beyond ±20 kb are drawn at the edge. Each panel is scaled to its own peak, so heights compare between the two tools within a panel and not between species. Gray band, ±50 nt; triangles, medians; the text in each panel gives the share of eligible transcripts with a UTR annotated by SQANTI-evidence and by BRAKER4. BRAKER4 3′ ends fall short of the poly(A) site in 49–81% of transcripts. SQANTI-evidence 5′ ends lie beyond the annotated start in 66–85% of transcripts in five species, because IsoQuant takes the furthest supported read start as the start of the model. BRAKER4 5′ ends lie beyond the annotated start in 22–81% of the transcripts for which it annotates a 5′ UTR.

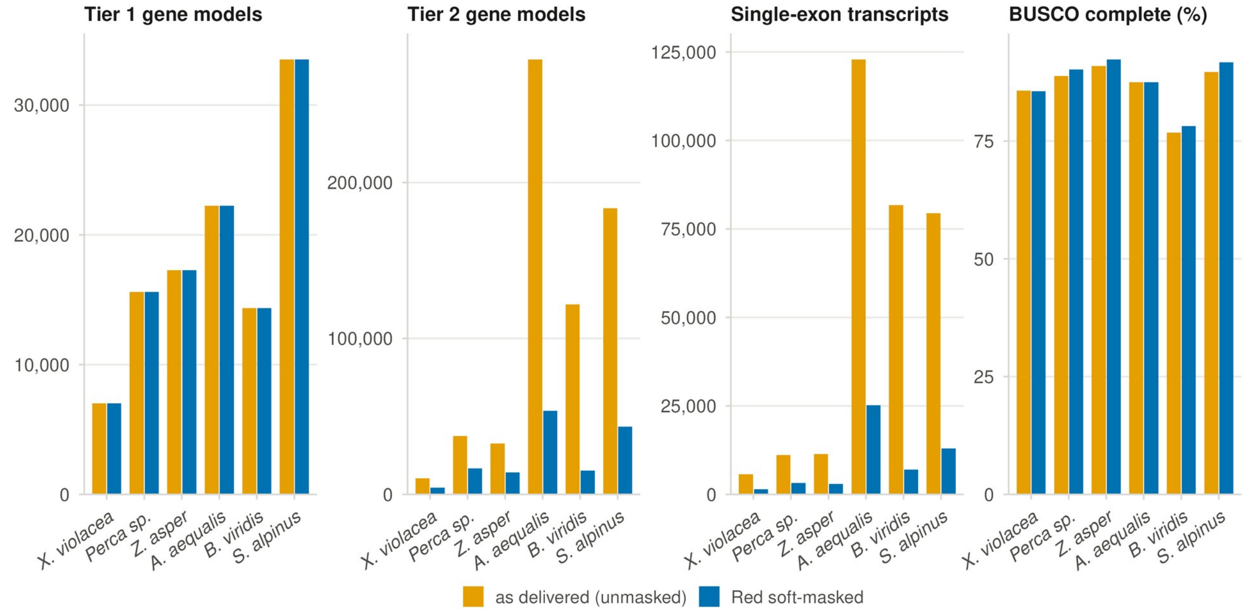


**Figure S15 | Soft-masking removes most of the predicted layer and leaves the curated layer unchanged.** The six pilot assemblies annotated as delivered (unmasked) and after Red soft-masking: Tier 1 gene models, Tier 2 gene models, single-exon transcripts and annotation BUSCO completeness. Masking removes 55–88% of the Tier 2 genes and 71–91% of the single-exon transcripts, while Tier 1 changes by at most four genes and completeness holds or rises in five of six genomes. *B. viridis* is scored on *tetrapoda_odb12*.

### Supplementary tables

| Species | Transcript reconstruction | Filter | Tier 1 transcripts | Tier 1, locus | Tier 1, transcript | Tier 2, locus | Tier 2, transcript | Whole annotation, locus | Whole annotation, transcript |
|---|---|---|---|---|---|---|---|---|---|
| *S. cerevisiae* | IsoQuant | Strict | 3,164 | 44.7 / 95.8 | 43.9 / 91.5 | 24.3 / 88.0 | 23.8 / 78.5 | 68.9 / 92.9 | 67.7 / 86.5 |
| | | Flexible | 4,045 | 55.5 / 94.8 | 54.6 / 89.0 | 14.1 / 85.7 | 13.9 / 76.0 | 69.7 / 92.8 | 68.4 / 86.0 |
| | | None | 4,304 | 57.9 / 94.2 | 56.9 / 87.2 | 12.3 / 85.2 | 12.1 / 75.0 | 70.2 / 92.5 | 69.0 / 84.8 |
| | IsoSeq3 | Strict | 20,501 | 68.5 / 93.7 | 67.4 / 21.7 | 6.7 / 87.0 | 6.6 / 75.8 | 75.2 / 93.0 | 74.0 / 23.2 |
| | | Flexible | 22,129 | 71.5 / 94.9 | 70.3 / 21.0 | 5.3 / 86.2 | 5.2 / 74.2 | 76.8 / 94.2 | 75.5 / 22.1 |
| | | None | 27,880 | 74.5 / 94.5 | 73.3 / 17.3 | 3.3 / 84.1 | 3.2 / 73.9 | 77.7 / 94.0 | 76.5 / 17.9 |
| *C. elegans* | IsoQuant | Strict | 12,124 | 42.4 / 87.6 | 31.8 / 74.3 | 17.5 / 46.2 | 12.4 / 43.9 | 60.0 / 69.4 | 44.2 / 62.2 |
| | | Flexible | 13,376 | 46.3 / 86.9 | 34.7 / 73.5 | 15.4 / 44.9 | 10.9 / 42.5 | 61.6 / 70.4 | 45.6 / 62.6 |
| | | None | 15,485 | 48.2 / 83.6 | 36.3 / 66.5 | 13.0 / 44.7 | 9.2 / 41.9 | 61.2 / 70.5 | 45.5 / 59.4 |
| | IsoSeq3 | Strict | 119,233 | 46.9 / 78.3 | 40.9 / 9.7 | 11.4 / 42.7 | 8.0 / 40.2 | 58.2 / 67.3 | 49.0 / 11.1 |
| | | Flexible | 129,422 | 48.7 / 79.7 | 42.7 / 9.3 | 10.9 / 42.4 | 7.7 / 39.9 | 59.6 / 68.6 | 50.4 / 10.6 |
| | | None | 160,877 | 50.9 / 78.1 | 45.1 / 7.9 | 9.4 / 42.3 | 6.6 / 39.7 | 60.3 / 69.0 | 51.7 / 8.8 |
| *A. thaliana* | IsoQuant | Strict | 14,529 | 42.2 / 97.5 | 29.9 / 82.8 | 19.3 / 53.7 | 13.2 / 49.7 | 61.4 / 77.6 | 43.1 / 68.8 |
| | | Flexible | 14,972 | 43.4 / 97.4 | 30.7 / 82.7 | 18.6 / 53.6 | 12.8 / 49.7 | 62.0 / 78.2 | 43.5 / 69.2 |
| | | None | 16,796 | 45.8 / 96.2 | 32.7 / 78.5 | 16.9 / 53.1 | 11.6 / 49.2 | 62.7 / 79.0 | 44.3 / 67.9 |
| | IsoSeq3 | Strict | 22,147 | 38.5 / 97.1 | 27.7 / 50.3 | 21.0 / 54.2 | 14.4 / 50.5 | 59.4 / 75.9 | 42.0 / 50.4 |
| | | Flexible | 23,225 | 39.5 / 96.6 | 28.4 / 49.3 | 20.4 / 54.1 | 14.0 / 50.3 | 59.8 / 76.2 | 42.4 / 49.7 |
| | | None | 27,950 | 42.4 / 94.9 | 30.7 / 44.3 | 18.6 / 54.1 | 12.7 / 50.3 | 60.9 / 77.2 | 43.5 / 45.9 |
| *D. melanogaster* | IsoQuant | Strict | 8,319 | 41.7 / 94.5 | 28.7 / 76.5 | 27.3 / 58.1 | 17.2 / 54.5 | 68.9 / 75.7 | 45.9 / 66.5 |
| | | Flexible | 8,519 | 42.1 / 94.1 | 29.1 / 75.6 | 26.9 / 58.0 | 17.0 / 54.4 | 69.0 / 75.7 | 46.0 / 66.1 |
| | | None | 8,896 | 42.7 / 93.8 | 29.6 / 73.9 | 25.6 / 57.6 | 16.1 / 53.8 | 68.3 / 75.9 | 45.7 / 65.3 |
| *G. gallus* | IsoQuant | Strict | 6,045 | 12.7 / 54.3 | 5.5 / 38.2 | 10.7 / 15.7 | 4.3 / 15.6 | 23.4 / 25.6 | 9.8 / 23.3 |
| | | Flexible | 6,268 | 13.2 / 54.7 | 5.7 / 38.4 | 10.7 / 15.7 | 4.3 / 15.6 | 23.9 / 26.0 | 10.0 / 23.6 |
| | | None | 7,298 | 14.1 / 52.9 | 6.1 / 35.2 | 10.3 / 15.8 | 4.1 / 15.6 | 24.4 / 26.5 | 10.2 / 23.3 |
| | IsoSeq3 | Strict | 13,427 | 13.0 / 55.9 | 5.9 / 18.7 | 10.9 / 15.9 | 4.4 / 15.8 | 23.8 / 26.0 | 10.3 / 17.4 |
| | | Flexible | 13,897 | 13.3 / 56.1 | 6.1 / 18.5 | 10.8 / 15.9 | 4.4 / 15.8 | 24.1 / 26.3 | 10.4 / 17.3 |
| | | None | 16,812 | 14.1 / 54.1 | 6.5 / 16.4 | 10.5 / 15.9 | 4.2 / 15.8 | 24.6 / 26.7 | 10.7 / 16.1 |
| *D. rerio* | IsoQuant | Strict | 18,799 | 34.3 / 62.0 | 21.9 / 48.6 | 14.8 / 14.7 | 9.1 / 14.2 | 49.0 / 31.4 | 31.0 / 28.4 |
| | | Flexible | 23,755 | 39.8 / 59.7 | 25.4 / 44.7 | 12.5 / 13.3 | 7.7 / 12.8 | 52.2 / 32.4 | 33.2 / 28.3 |
| | | None | 40,192 | 45.5 / 50.9 | 29.5 / 30.6 | 8.3 / 9.7 | 5.2 / 9.2 | 53.8 / 30.6 | 34.6 / 22.7 |
| | IsoSeq3 | Strict | 65,949 | 34.8 / 61.5 | 23.0 / 14.6 | 13.9 / 14.9 | 8.6 / 14.4 | 48.6 / 32.3 | 31.6 / 14.5 |
| | | Flexible | 72,882 | 37.6 / 61.6 | 25.0 / 14.3 | 12.8 / 14.3 | 7.9 / 13.8 | 50.4 / 33.4 | 33.0 / 14.2 |
| | | None | 131,051 | 43.7 / 52.7 | 29.7 / 9.5 | 9.2 / 12.0 | 5.7 / 11.5 | 52.9 / 33.0 | 35.5 / 9.8 |
| *M. musculus* | IsoQuant | Strict | 28,444 | 52.2 / 81.7 | 24.1 / 48.0 | 7.7 / 16.3 | 3.0 / 15.5 | 59.9 / 53.9 | 27.1 / 38.9 |
| | | Flexible | 29,412 | 53.6 / 81.6 | 24.8 / 47.8 | 7.4 / 16.1 | 2.9 / 15.3 | 61.0 / 54.6 | 27.7 / 39.1 |
| | | None | 34,312 | 55.1 / 75.7 | 25.8 / 42.7 | 6.4 / 14.5 | 2.5 / 13.7 | 61.5 / 52.5 | 28.3 / 36.0 |
| | IsoSeq3 | Strict | 78,254 | 36.3 / 77.8 | 18.8 / 13.6 | 12.1 / 22.1 | 4.7 / 21.5 | 48.4 / 47.7 | 23.5 / 14.7 |
| | | Flexible | 79,890 | 36.8 / 78.1 | 19.1 / 13.5 | 12.0 / 22.0 | 4.7 / 21.3 | 48.8 / 47.9 | 23.7 / 14.6 |
| | | None | 96,273 | 38.3 / 72.5 | 20.2 / 11.9 | 10.7 / 20.9 | 4.2 / 20.3 | 49.0 / 47.1 | 24.4 / 12.8 |
| *H. sapiens* | IsoQuant | Strict | 30,161 | 52.3 / 81.4 | 9.7 / 50.1 | 11.1 / 15.4 | 1.4 / 14.6 | 63.2 / 46.6 | 11.1 / 38.4 |
| | | Flexible | 31,563 | 54.5 / 81.5 | 10.2 / 50.2 | 10.5 / 14.9 | 1.3 / 14.1 | 64.9 / 47.3 | 11.5 / 38.7 |
| | | None | 37,286 | 56.5 / 78.5 | 10.8 / 45.1 | 9.4 / 13.9 | 1.2 / 13.1 | 65.8 / 47.3 | 12.0 / 36.3 |
| | IsoSeq3 | Strict | 154,692 | 58.3 / 82.1 | 16.1 / 16.2 | 9.0 / 13.8 | 1.1 / 13.1 | 67.2 / 49.5 | 17.2 / 16.0 |

| Species | Transcript reconstruction | Filter | Tier 1 transcripts | Tier 1, locus | Tier 1, transcript | Tier 2, locus | Tier 2, transcript | Whole annotation, locus | Whole annotation, transcript |
|---|---|---|---|---|---|---|---|---|---|
| | | Flexible | 157,866 | 59.3 / 81.8 | 16.4 / 16.2 | 8.8 / 13.4 | 1.1 / 12.7 | 68.0 / 49.4 | 17.5 / 15.9 |
| | | None | 194,378 | 60.9 / 74.4 | 17.7 / 14.2 | 7.9 / 12.0 | 1.0 / 11.2 | 68.7 / 46.6 | 18.7 / 14.0 |

**Table S1 | Per-tier CDS accuracy of every SQANTI-evidence run.** Sensitivity / precision (%) at the CDS locus level and at the transcript level (exact CDS intron chain) of Tier 1 alone, Tier 2 alone and the whole annotation, for the IsoQuant and IsoSeq3 reconstructions with the strict, flexible and no filter, eight species (the Nanopore *D. melanogaster* dataset has no IsoSeq3 run). Tier 1 transcripts, number of curated transcripts the run retained.

| Hint feature set | Human chr19, locus F1 | Human chr19, transcript F1 | *Arabidopsis*, locus F1 | *Arabidopsis*, transcript F1 |
|---|---|---|---|---|
| intron + CDS | 59.8 | 15.8 | 69.0 | 54.2 |
| intron + CDS + start + stop (default) | 59.5 | 15.7 | 69.0 | 54.2 |
| CDS only | 59.0 | 15.6 | 68.9 | 54.1 |
| intron only | 57.7 | 14.6 | 68.7 | 53.6 |
| CDS + exon | 57.2 | 15.3 | 68.7 | 53.2 |
| all six feature types | 54.8 | 15.2 | 68.5 | 53.1 |
| no long-read hints (protein hints only) | 53.0 | 13.3 | 67.5 | 52.7 |
| exon only | 51.3 | 13.1 | 68.1 | 52.5 |
| intron + exon | 50.0 | 12.9 | 66.5 | 51.2 |
| **Weights (default feature set)** | **Human chr19, locus F1** | **Human chr19, transcript F1** | ***Arabidopsis*, locus F1** | ***Arabidopsis*, transcript F1** |
| default (bonus saturating, malus 1) | 59.5 | 15.7 | 69.0 | 54.2 |
| long-read bonus $10^{15}$ | 59.3 | 15.7 | 68.9 | 54.1 |
| long-read bonus $10^4$ | 59.2 | 15.5 | 68.9 | 54.0 |
| malus 0.9 | 43.2 | 9.2 | 31.9 | 23.7 |
| malus 0.5 | 2.1 | 0.4 | 0.0 | 0.0 |
| malus 0.1 | 0.0 | 0.0 | 0.0 | 0.0 |

**Table S2 | Hint feature sets and weights: locus and transcript-level F1 of the raw AUGUSTUS prediction.** Human chromosome 19 and whole-genome *Arabidopsis*, trained model fixed, scored against the reference CDS before the two layers were combined. Top, hint feature sets; bottom, weights with the default feature set. The anchor rerun of the production configuration reproduced the production prediction; differences below 0.5 F1 are within the noise of alternative-transcript ordering.

| Species | Transcripts | Exact (%) | Reference transcripts exact | Reference transcripts found by one set only | Reference genes found by one set only |
|---|---|---|---|---|---|
| *S. cerevisiae* | 4,045 / 4,079 | 92.9 / 94.0 | 3,601 / 3,818 | 14 / 231 | 14 / 231 |
| *C. elegans* | 13,376 / 13,181 | 77.0 / 70.9 | 9,832 / 9,285 | 867 / 320 | 437 / 232 |
| *A. thaliana* | 14,972 / 14,138 | 93.0 / 88.3 | 12,372 / 12,360 | 213 / 201 | 213 / 201 |
| *D. melanogaster* | 8,519 / 7,298 | 87.5 / 82.1 | 6,427 / 5,983 | 600 / 156 | 199 / 96 |
| *G. gallus* | 6,268 / 5,450 | 45.0 / 42.4 | 2,401 / 2,296 | 159 / 54 | 72 / 36 |
| *D. rerio* | 23,755 / 22,927 | 49.3 / 45.1 | 10,586 / 10,224 | 771 / 409 | 439 / 288 |
| *M. musculus* | 29,412 / 20,485 | 62.4 / 59.7 | 13,985 / 12,117 | 2,194 / 326 | 636 / 183 |
| *H. sapiens* | 31,563 / 20,689 | 63.8 / 59.3 | 15,782 / 12,166 | 4,105 / 489 | 488 / 154 |

**Table S3 | Tier 1 against the AUGUSTUS models it replaces, at the same loci.** Flexible-filter run, eight species; each cell gives Tier 1 / AUGUSTUS. Transcripts: Tier 1, every curated model of the annotation; AUGUSTUS, the raw AUGUSTUS prediction, before the two layers were combined, of every AUGUSTUS gene that overlaps a Tier 1 gene on the same strand, that is, the models discarded when the two layers are combined. Both sets are scored with gffcompare on the CDS against the reference of the benchmark; exact, gffcompare class =, as a percentage of the models of the set (per-model exact-CDS precision, as in Fig. S10). Reference transcripts exact, distinct reference transcripts matched exactly by at least one model of the set; found by one set only, those matched by that set and not the other, counted by reference transcript and by reference gene. Tier 1 is exact more often and recovers more reference transcripts in seven of eight species; *S. cerevisiae* is the exception.

| Species | AUGUSTUS models | Discarded (n, %) | Intron chain already in Tier 1 (n, % of discarded) | New multi-exon, reference-exact | New multi-exon, non-exact | Spurious per exact | Single-exon, exact / total |
|---|---|---|---|---|---|---|---|
| *S. cerevisiae* | 5,300 | 4,094 (77.2%) | 3,575 (87.3%) | 0 | 52 | n/a | 338 / 467 |
| *C. elegans* | 20,552 | 13,301 (64.7%) | 10,728 (80.7%) | 306 | 2,078 | 6.8 | 0 / 189 |
| *A. thaliana* | 24,731 | 14,355 (58.0%) | 12,943 (90.2%) | 157 | 893 | 5.7 | 138 / 362 |
| *D. melanogaster* | 14,367 | 7,483 (52.1%) | 6,262 (83.7%) | 99 | 810 | 8.2 | 0 / 312 |
| *G. gallus* | 17,148 | 5,588 (32.6%) | 4,664 (83.5%) | 50 | 714 | 14.3 | 0 / 160 |
| *D. rerio* | 40,285 | 20,913 (51.9%) | 14,444 (69.1%) | 383 | 3,935 | 10.3 | 0 / 2,151 |
| *M. musculus* | 31,810 | 21,144 (66.5%) | 16,185 (76.5%) | 282 | 3,448 | 12.2 | 0 / 1,229 |
| *H. sapiens* | 31,450 | 19,518 (62.1%) | 15,109 (77.4%) | 439 | 3,085 | 7.0 | 0 / 885 |

**Table S4 | AUGUSTUS models discarded by the combination rule.** AUGUSTUS models of the flexible-filter run that were discarded because they overlapped a curated gene on the same strand, eight species. Discarded, number and percentage of the AUGUSTUS models; intron chain already in Tier 1, discarded models whose CDS intron chain a curated transcript already contained (percentage of the discarded models); new multi-exon, discarded multi-exon models with a CDS intron chain absent from Tier 1, split by whether a reference transcript carries that chain (reference-exact) or not (non-exact); spurious per exact, the ratio of the two (n/a in *S. cerevisiae*, which has no reference-exact multi-exon discards); single-exon, discarded single-exon models whose CDS matches a reference transcript, over all discarded single-exon models. Across the eight species, 1,716 new multi-exon models were reference-exact and 15,015 were not.

| Species | A | B | C | D | E | BRAKER3 | BRAKER4 | Tier 1 loci overlapping a coding gene (%) | BRAKER3 models overlapping a coding gene (%) | BRAKER4 models overlapping a coding gene (%) |
|---|---|---|---|---|---|---|---|---|---|---|
| *C. elegans* | 61.6 / 72.0 | 47.8 / 84.7 | 46.9 / 86.8 | 46.3 / 87.3 | 46.3 / 87.6 | 66.1 / 84.4 | 81.4 / 90.0 | 97.8 | 94.0 | 95.2 |
| *A. thaliana* | 62.0 / 78.6 | 52.0 / 92.7 | 45.9 / 96.8 | 43.4 / 97.4 | 43.4 / 97.8 | 69.1 / 92.0 | 80.0 / 91.7 | 99.5 | 97.2 | 96.6 |
| *D. melanogaster* | 69.0 / 79.5 | 53.6 / 90.4 | 42.3 / 94.2 | 42.1 / 94.4 | 42.1 / 94.5 | 68.4 / 89.3 | 86.1 / 86.3 | 99.5 | 97.1 | 94.4 |
| *G. gallus* | 23.9 / 27.9 | 16.6 / 49.1 | 13.7 / 54.3 | 13.2 / 55.2 | 13.2 / 55.7 | 19.4 / 52.3 | 48.3 / 52.5 | 98.2 | 93.7 | 91.1 |
| *D. rerio* | 52.2 / 33.7 | 45.1 / 46.6 | 42.8 / 57.1 | 39.7 / 60.3 | 39.7 / 75.0 | 57.2 / 61.2 | 66.3 / 57.9 | 78.7 | 75.6 | 73.9 |
| *M. musculus* | 60.9 / 56.0 | 57.6 / 66.0 | 54.1 / 81.2 | 53.6 / 82.2 | 53.6 / 83.4 | 65.6 / 78.6 | 76.9 / 76.6 | 95.1 | 88.6 | 86.2 |
| *H. sapiens* | 64.8 / 48.5 | 60.1 / 62.3 | 55.5 / 77.5 | 54.4 / 81.8 | 54.4 / 88.5 | 64.3 / 70.9 | 80.3 / 64.4 | 91.3 | 80.9 | 77.4 |

**Table S5 | Subsets of the SQANTI-evidence annotation, seven species.** CDS locus sensitivity / precision (%) of the subsets defined in Methods: A, default annotation; B, Tier 2 restricted to models with every intron protein-supported; C, B without single-exon Tier 2 models; D, Tier 1 alone; E, D with Tier 1 loci that overlap no annotated gene excluded from the precision denominator. BRAKER annotations as in Fig. 3. The three right-hand columns give the percentage of Tier 1 loci, BRAKER3 models and BRAKER4 models that overlap an annotated coding gene on the same strand, whatever their structure. Scenario values are recomputed from the gffcompare tracking files; they differ from the whole-annotation values of Fig. 3, which come from the gffcompare summary, by up to 3.8 points of precision (*D. melanogaster*) and 0.1 points of sensitivity, and from the Tier 1 values of Table S1 by up to 0.6 points of precision.

| Species | Genes | | Sensitivity, genes with read evidence (%) | | | Sensitivity, genes without read evidence (%) | | |
|---|---|---|---|---|---|---|---|---|
| | With read evidence, n (%) | Without, n (%) | SQANTI-evidence | BRAKER3 | BRAKER4 | SQANTI-evidence | BRAKER3 | BRAKER4 |
| *C. elegans* | 11,891 (59.5%) | 8,094 (40.5%) | 82.3 | 87.7 | 92.3 | 31.2 | 34.4 | 65.2 |
| *A. thaliana* | 13,363 (48.4%) | 14,265 (51.6%) | 93.5 | 96.3 | 96.5 | 32.6 | 43.6 | 64.4 |
| *D. melanogaster* | 6,654 (47.6%) | 7,332 (52.4%) | 90.8 | 93.6 | 96.2 | 48.9 | 45.3 | 76.6 |
| *G. gallus* | 5,274 (31.0%) | 11,733 (69.0%) | 45.4 | 50.4 | 62.5 | 14.2 | 5.5 | 41.8 |
| *D. rerio* | 17,586 (68.7%) | 8,006 (31.3%) | 64.7 | 71.9 | 75.5 | 24.2 | 24.1 | 45.1 |
| *M. musculus* | 14,496 (65.0%) | 7,805 (35.0%) | 83.5 | 87.4 | 89.9 | 17.6 | 23.5 | 50.6 |
| *H. sapiens* | 12,823 (62.5%) | 7,710 (37.5%) | 85.8 | 87.6 | 88.9 | 23.7 | 19.8 | 58.7 |

**Table S6 | CDS locus sensitivity at reference coding genes with and without read evidence.** Genes with read evidence, reference coding genes that at least one IsoQuant model overlaps before filtering, as a count and a percentage of the coding genes; genes without, the remainder. Sensitivity, percentage of the genes of each stratum for which at least one predicted transcript reproduces the CDS intron chain exactly (gffcompare class =). Genes are counted per reference gene, not per gffcompare locus as in Table S1. SQANTI-evidence with the flexible filter, BRAKER3 run with the same reads, and BRAKER4, seven species.

| Species | Tier 2 models | Reference-exact, n (%) | Non-exact, n | Non-exact models by what they overlap, n (% of non-exact) | | | |
|---|---|---|---|---|---|---|---|
| | | | | Non-coding gene or pseudogene | Coding gene, different structure | No annotated gene, unmasked | No annotated gene, mostly soft-masked |
| *C. elegans* | 7,251 | 3,086 (42.6%) | 4,165 | 678 (16.3%) | 3,222 (77.4%) | 179 (4.3%) | 86 (2.1%) |
| *A. thaliana* | 10,376 | 5,188 (50.0%) | 5,188 | 38 (0.7%) | 4,594 (88.6%) | 359 (6.9%) | 197 (3.8%) |
| *D. melanogaster* | 6,919 | 3,777 (54.6%) | 3,142 | 104 (3.3%) | 2,881 (91.7%) | 130 (4.1%) | 27 (0.9%) |
| *G. gallus* | 11,631 | 1,815 (15.6%) | 9,816 | 234 (2.4%) | 8,465 (86.2%) | 976 (9.9%) | 141 (1.4%) |

| Species | Tier 2 models | Reference-exact, n (%) | Non-exact, n | Non-exact models by what they overlap, n (% of non-exact) | | | |
|---|---|---|---|---|---|---|---|
| | | | | Non-coding gene or pseudogene | Coding gene, different structure | No annotated gene, unmasked | No annotated gene, mostly soft-masked |
| *D. rerio* | 25,200 | 3,232 (12.8%) | 21,968 | 785 (3.6%) | 7,953 (36.2%) | 4,555 (20.7%) | 8,675 (39.5%) |
| *M. musculus* | 10,677 | 1,636 (15.3%) | 9,041 | 3,860 (42.7%) | 3,315 (36.7%) | 758 (8.4%) | 1,108 (12.3%) |
| *H. sapiens* | 14,796 | 2,094 (14.2%) | 12,702 | 4,456 (35.1%) | 4,218 (33.2%) | 3,031 (23.9%) | 997 (7.8%) |

**Table S7 | Tier 2 models by genomic context.** Predicted-layer models of the flexible-filter run, seven species. Reference-exact, models whose CDS intron chain matches a reference transcript (percentage of the Tier 2 models); the non-exact models are split by what they overlap on the same strand, with percentages of the non-exact models: a non-coding gene or pseudogene only (absent from the CDS reference), a coding gene with a different structure, no annotated gene in unmasked sequence, or no annotated gene in sequence more than 50% soft-masked.

| Species | BUSCO lineage | Tier 1 genes | | Tier 2 genes | | Single-exon transcripts | | BUSCO C (%) | |
|---|---|---|---|---|---|---|---|---|---|
| | | Unmasked | Red | Unmasked | Red | Unmasked | Red | Unmasked | Red |
| *Xylocopa violacea* | apoidea_odb12 | 7,024 | 7,025 | 10,236 | 4,391 | 5,692 | 1,440 | 85.7 | 85.6 |
| *Perca* sp. ‘yellow fin Alpine’ | actinopterygii_odb12 | 15,604 | 15,604 | 37,479 | 16,772 | 11,137 | 3,210 | 88.8 | 90.2 |
| *Zingel asper* | actinopterygii_odb12 | 17,271 | 17,269 | 32,644 | 14,109 | 11,430 | 2,945 | 90.9 | 92.3 |
| *Alopecurus aequalis* | poales_odb12 | 22,242 | 22,246 | 278,817 | 53,694 | 122,844 | 25,233 | 87.5 | 87.5 |
| *Bufotes viridis* | tetrapoda_odb12 | 14,354 | 14,354 | 121,845 | 15,216 | 81,754 | 7,009 | 76.8 | 78.2 |
| *Salvelinus alpinus* | actinopterygii_odb12 | 33,505 | 33,504 | 183,588 | 43,381 | 79,434 | 12,969 | 89.7 | 91.7 |

**Table S8 | Effect of Red soft-masking on the six ERGA pilot genomes.** Each genome was annotated twice with the same reads and settings, first as delivered (unmasked) and then after soft-masking with Red [43]. Unmasked, the genome as delivered; Red, after soft-masking. Tier 1 genes, curated genes; Tier 2 genes, AUGUSTUS genes kept when the two layers are combined; single-exon transcripts, whole annotation; BUSCO C, annotation BUSCO completeness from GAQET2 on the lineage in the second column. Masking removes 55–88% of the Tier 2 genes and changes Tier 1 by at most four genes.

#### *Supplementary notes*

**Note S1. Read-quality audit.** *C. elegans* initially scored implausibly low (locus sensitivity 40.7%), and the alignment error rate of its reads was 11.9%; yeast was at 12.7% and chicken at 14.1%, whereas HiFi reads are below 2%. The three SRA runs had been downloaded as FASTQ, which strips the PacBio read names, and wrapped into CCS-style BAMs by a conversion script, so that lima and *isoseq refine* accepted subreads as if they were HiFi reads. The worm dataset was replaced with N2 Revio Kinnex FLNC data (PRJEB121150, three replicates), the yeast dataset with four pooled Sequel II runs (PRJNA883530), and the chicken dataset by regenerating HiFi reads with *ccs* from the submitted subreads BAM of ERR7015103, followed by lima and *isoseq refine*. The pipeline now reports the alignment error rate of the reads before it starts.

**Note S2. Rejected filter alternatives.** A single-exon guard, a relaxed intra-priming check applied only to single-exon models, removed 145 reference-exact isoforms in yeast, where 91% of transcripts are single-exon, while keeping all 17 intergenic or intronic models it was meant to exclude, and was rejected. A two-branch design that would select the rule set according to the measured noise of the library was not pursued; instead, the workflow reports the noise diagnostic before filtering (see Results), so that the user can choose the strict filter for a noisy library.

**Note S3. Selecting Tier 2 models by protein support after the run.** The tier of every model is written in the annotation (*tier=SQANTI_curated* for Tier 1 and *tier=Augustus* for Tier 2, on every feature), so Tier 1 alone or the whole annotation is selected with a single filter on that attribute. Selecting Tier 2 models by protein support, as in steps B and C of Fig. 5, also needs the protein support of each intron, which the workflow does not write to the annotation. It is computed from the protein-hint file that every run writes (evidence_driven/hints/<species>.protein.hints.gff) by matching each intron of a Tier 2 model to the protein-hint introns, with the script of the benchmark repository that produced Fig. 5 (bin/comparison/braker_compare.py; see Availability of data and materials). Step E requires the reference annotation.
